# Resolving the dynamics of photosynthetically produced ROS by high-resolution monitoring of chloroplastic E_GSH_ in *Arabidopsis*

**DOI:** 10.1101/2020.03.04.976092

**Authors:** Zachary Haber, Shilo Rosenwasser

**Author notes:** Corresponding author. (S.R.).

## Abstract

Plants are naturally subjected to fluctuations in light intensity, causing unbalanced photosynthetic electron fluxes and overproduction of reactive oxygen species (ROS). While high rates of ROS production are harmful, moderate levels play a signaling role, coordinating photosynthetic activity and downstream metabolism. Here, we explore the dynamics of light-dependent oxidant production by high-temporal-resolution monitoring of chloroplastic glutathione redox potential (chl-E_GSH_) using chloroplast-targeted-roGFP2-expressing *Arabidopsis* lines, over several days, under dynamic environmental conditions and in correlation with PSII operating efficiency. Peaks in chl-E_GSH_ oxidation during light-darkness transitions, when light harvesting is not balanced with downstream metabolism, were observed. Increasing light intensities triggered a binary oxidation response, with a threshold around the light saturating point, pointing for two regulated oxidative states of the chl-E_GSH_. These patterns were not affected in *npq1* plants which are impaired in non-photochemical quenching. Frequency-dependent oscillations between the two oxidation states were observed under fluctuating light in WT and *npq1* plants, but not in *pgr5* plants, suggesting a role for PSI photoinhibition in regulation of oxidant production. Remarkably, *pgr5* plants showed an increase in chl-E_GSH_ oxidation during the nights following light stresses, linking between day photoinhibition and night redox metabolism. This work provides a comprehensive view on the balance between photosynthesis-dependent ROS production and antioxidant activity during light acclimation.

## Introduction

As sessile organisms that grow under highly variable light intensities, plants must constantly sense, respond, and adapt to the instantaneous modulation of daytime photon fluxes. This regulation is critical when absorption of light energy, and consequent production of NADPH and ATP, exceed the capacity of downstream reactions, such as during light-darkness transitions or under high or fluctuating light (Kono and Terashima, 2014; Tikhonov, 2015), as over-reduction of the electron transport chain can lead to the production of deleterious reactive oxygen species (ROS).

Several mechanisms have evolved in plants to achieve homeostasis and optimal photosynthetic performance under fluctuating environmental conditions, thereby protecting the photosynthesis machinery from light stresses (photoprotection). Among them are energy dissipation via heat, known as non-photochemical quenching (NPQ; Li et al., 2000), cyclic electron flow (CEF; Munekage et al., 2002) and the water-water cycle (WWC; Mehler, 1951). Plants also cope with ROS overproduction by generating small molecular antioxidants such as glutathione and ascorbate, and enzymatic antioxidants such as ascorbate peroxidase (APX) and peroxiredoxins (Mittler et al., 2004; Takahashi and Badger, 2011; Kono and Terashima, 2014).

The rapid NPQ component, known as qE, requires the activity of violaxanthin de-epoxidase (VDE), which catalyzes the conversion of violaxanthin into zeaxanthin (Niyogi et al., 1998). Analysis of the VDE null mutant (*npq1*), demonstrated that this mechanism increases plant tolerance to variation in light intensity by protecting photosystem II (PSII) under field and fluctuating light conditions (Külheim et al., 2002).

CEF plays a role in balancing the stromal ATP/NADPH ratio in accordance with their demand for primary metabolism and for protection of photosystem I (PSI) against photoinhibition (Avenson et al., 2005; Shikanai, 2007; Takahashi et al., 2009), thereby serving as an electron sink under excessive light conditions (Asada, 1999; Ort and Baker, 2002). In higher plants, two CEF pathways are known to exist: the PROTON GRADIENT REGULATION 5 (PGR5)-dependent pathway, which is believed to be the main pathway in C3 plants, and the NADH dehydrogenase-like (NDH)-complex-dependent pathway (Shikanai, 2016). The PGR5 protein has been shown to be a key player in protecting PSI functionality (Munekage et al., 2002; Suorsa et al., 2012; Suorsa et al., 2013) via regulation of CEF, though the mechanism is not yet clear. The *Arabidopsis* PGR5 null mutant *(pgr5)* showed stunted growth, impaired NPQ and lower electron transport rate (ETR), relative to wild type (WT), under high-light and fluctuating light, but not under normal growth light conditions (Munekage et al., 2002). Therefore, exposure of *pgr5* to increased light intensities has served as an experimental model for assessing the physiological effects of PSI photoinhibition (Tiwari et al., 2016).

During WWC, electrons are donated from PSI to molecular oxygen in the Mehler reaction, yielding superoxide radicals (O_2_·^−^), which are then dismutated to molecular oxygen (O_2_) and H_2_O_2_ in a reaction catalyzed by superoxide dismutase (SOD; Mehler, 1951; Rizhsky et al., 2003). Chloroplastic peroxidases involved in the detoxification of H_2_O_2_ include ascorbate peroxidases (APXs), glutathione peroxidases (GPXs) and peroxiredoxins (PRX; Awad et al., 2015). Reduced glutathione (GSH) is the major electron source for APX activity, and is supplied through the ascorbate-glutathione cycle (Foyer and Noctor, 2011), though ascorbate can also be regenerated by NADPH or ferredoxin (Asada, 1999). Thus, electrons for the detoxification of WWC-derived H_2_O_2_ are drawn, at least partially, from the cellular glutathione pool. Importantly, production of H_2_O_2_ through the WWC can play a regulatory role, allowing communication between the photosynthetic light reactions and downstream metabolic processes, by transmitting oxidative signals to redox-regulated proteins (Dangoor et al., 2012; Eliyahu et al., 2015).

In the past fifteen years, redox-sensitive green fluorescent proteins (roGFP) have allowed for *in vivo* monitoring of the glutathione redox state (E_GSH_) at high spatiotemporal resolution (Dooley et al., 2004; Hanson et al., 2004; Meyer et al., 2007; Gutscher et al., 2008; Meyer and Dick, 2010; Rosenwasser et al., 2010; Albrecht et al., 2011; Schwarzländer et al., 2015; Nietzel et al., 2019). Due to their high sensitivity, reversibility and insensitivity to pH alterations in the physiological range (Schwarzländer et al., 2008), such protein-based redox sensors are powerful tools for investigating redox dynamics in subcellular compartments. Recently, roGFP-expressing plants and algae have been tested under various biotic and abiotic stresses and had shown organelle-specific roGFP oxidation patterns (Rosenwasser et al., 2010; van Creveld et al., 2015; Bratt et al., 2016; Volpert et al., 2018; Mizrachi et al., 2019; Nietzel et al., 2019).

Here, we examined daily fluctuations in photosynthesis-dependent oxidants production under different light conditions, by developing an automatic system for monitoring the roGFP oxidation state throughout the day. We revealed unique patterns of chloroplast-specific E_GSH_ under normal, high-light and fluctuating light conditions. Moreover, examination of these patterns in *Arabidopsis* lines, mutated in key proteins involved in photoprotective mechanisms, uncovered their effect in controlling light-dependent ROS production.

## Results

### The chloroplastic E_GSH_ redox state is directly affected by photosynthetic electron transport

In order to map the *in vivo* temporal alterations in the chloroplastic E_GSH_ (chl-E_GSH_) under ‘steady state’ and light treatment conditions, we used a chloroplast-targeted redox-sensitive GFP 2 sensor (chl-roGFP2; Fig 1). Chloroplast targeting was achieved by using either the Transketolase signal peptide (chl-TKTP-roGFP2, 33) or a 2-Cys Peroxiredoxin A signal peptide (chl-PRXaTP-roGFP2, this study), as both target proteins to the chloroplast stroma (König et al., 2002; Schwarzländer et al., 2008). Chloroplast targeting was verified by the overlap of the chl-roGFP2 fluorescence signal with the chlorophyll fluorescence signal, as observed using confocal fluorescence microscopy (Fig. 1A). Whole-plant fluorescence images showed that fluorescence intensity following excitation at 405nm (emission at 510nm) increased and decreased following H_2_O_2_ and DTT applications, respectively; a contrasting pattern was observed following excitation at 465nm (Fig. 1B). Accordingly, ratiometric images (405nm/465nm) showed oxidation and reduction of the probe consequent to application of H_2_O_2_ and DTT, respectively (Fig. 1B), as well as a positive correlation between the concentration of H_2_O_2_ applied and chl-roGFP2 oxidation (Supp. Fig. 1). These results validated the sensitivity of chl-roGFP2 to redox alterations and are consistent with probe performance, previously observed in plant and mammalian cells (Dooley et al., 2004; Jiang et al., 2006; Schwarzländer et al., 2008).

**Figure 1:**
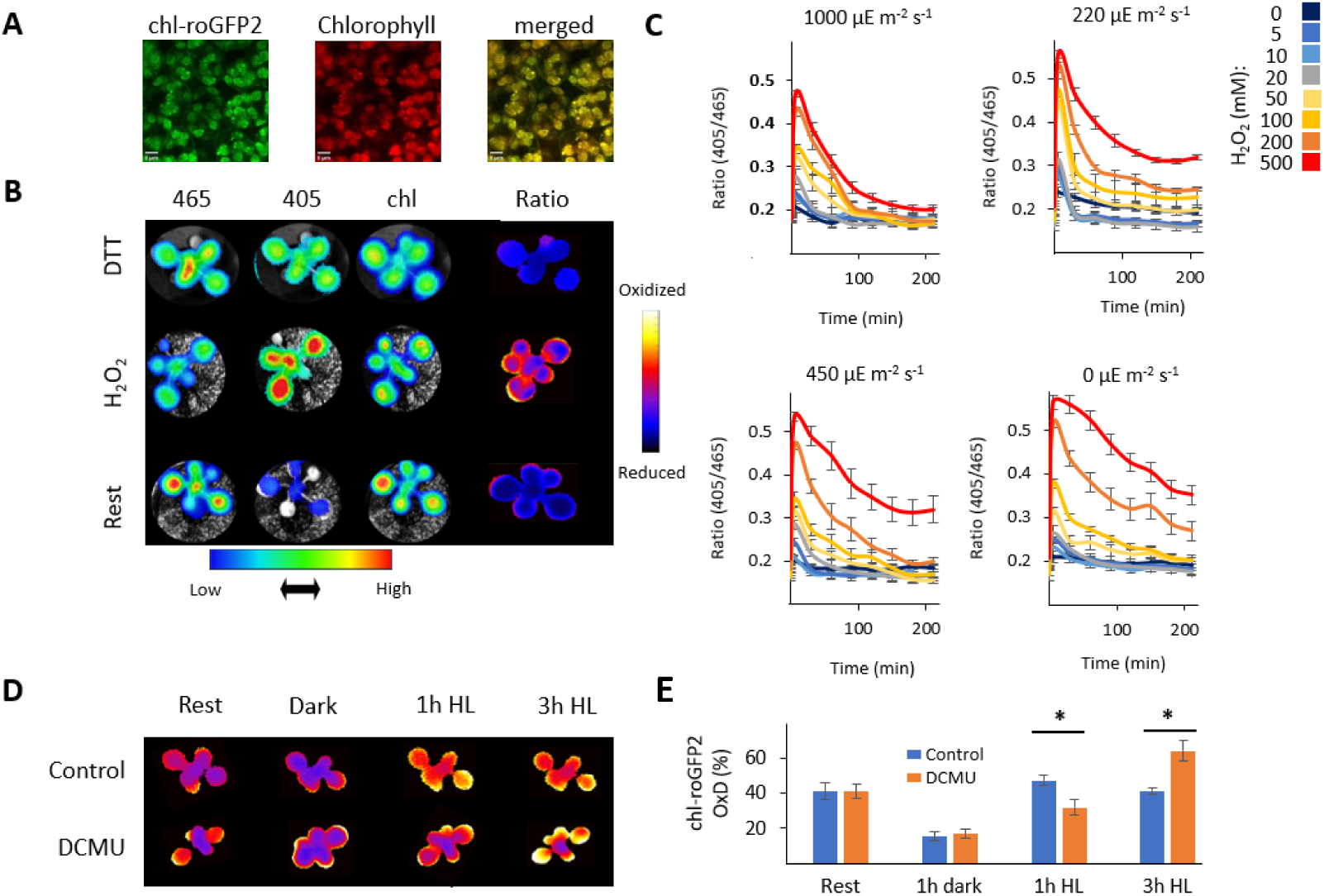
Light-dependent redox modification of chloroplast-targeted roGFP2. (A) Confocal images of *Arabidopsis* plants were acquired at 520nm, following excitation at 488nm for chl-roGFP2 fluorescence, and at 670nm, following excitation at 488nm for chlorophyll fluorescence. (B) Ratiometric analysis of chl-roGFP2 during rest and under fully oxidized and fully reduced conditions using whole-plant imaging. Emission was recorded at 510nm consequent to excitation at 405nm and 465nm. Chlorophyll fluorescence was detected by recording emission at 670nm following excitation at 405nm. Ratiometric images were generated by pixel-to-pixel division of the emission values at 510nm consequent to excitation at 405nm and 465nm. (C) The effect of light intensities on chl-roGFP2 reduction after application of H_2_O_2_, as examined by whole-plant imaging. *Arabidopsis* plants were treated with 0,5,10,20,50,100,200 or 500mM H_2_O_2_ and placed under light regimes of 0,220,450 or 1000µE m^-2^ s^-1^. Values represent means (of 6 plants) ± SE. (D&E) The effect of DCMU on chl-roGFP2 oxidation using whole-plant imaging and quantitative fluorometry. *Arabidopsis* plants were treated with 150µM DCMU or water (control) and placed under 700 µE m^-2^ s^-1^. chl-roGFP2 fluorescence ratios (405nm/465nm) were recorded before DCMU treatment, after 1 hour in the dark, and at 1- and 3-hours post-illumination. chl-roGFP2 oxidation was detected using whole-plant imaging (D) and as a function of time (minutes) using the fluorometer method (E). chl-roGFP oxidation degree values are presented as means (of 7-8 plants) ± SE. Asterisks (*) mark significant differences between the ratio between the control and the DCMU treatments (t-test, P<0.05).

To examine the interaction between the redox state of chl-roGFP2 and photosynthetic activity, we treated plants with various concentrations of H_2_O_2_ and measured the recovery of chl-roGFP2 from oxidation under different light intensities. chl-roGFP2 fluorescence was measured in Following H_2_O_2_ treatment, light-dependent reduction of the probe was fully achieved in plants treated with 500mM H_2_O_2_, only under the highest light intensity tested (1000µE m^-2^ s^-1^). Accordingly, the lowest reduction rates were observed in plants that were kept in the dark (Fig. 1C). In addition, probe oxidation was observed in plants exposed to one hour of high light conditions (700 µE m^-2^ s^-1^), but not in plants pretreated with 150µM 3-(3,4-dichlorophenyl)-1,1-dimethylurea (DCMU), as determined by whole-plant imaging (Fig. 1D) and quantitative fluorometry (Fig. 1E). Higher oxidation, compared to control, was observed in DCMU-treated plants after 3h under high-light (HL). Taken together, the chl-roGFP2 redox state is directly regulated by ROS production and by the reducing power produced in the photosynthetic electron transport.

### Distinct patterns of chloroplastic E_GSH_ during a diurnal cycle and in response to high-light conditions

The observed relationship between photosynthesis activity and chl-roGFP2 redox state, motivated us to systematically monitor its oxidation patterns under normal growth light and under changing light conditions. To this end, we developed an automated system that allows for continuous measurement of chl-roGFP2 oxidation degree (OxD) and chlorophyll fluorescence-derived photosynthetic parameters from many plants, under dynamic environmental conditions (Fig. 2). Examination of chl-roGFP2 oxidation under typical laboratory growth conditions, over the entire day (a 24h cycle of 16h 120 µE m^-2^ s^-1^/8h 0 µE m^-2^ s^-1^), showed a stable chl-roGFP2 state during the day, and an initial reduction (ΔOxD= −13%, compared to day ‘steady state’ conditions), followed by gradual oxidation during the night, nearly reaching the ‘steady state’ by the end of night (Fig. 2A). Interestingly, short peaks in chl-roGFP2 oxidation (ΔOxD= ∼ 10%) were observed during light-darkness transitions and vice versa (Fig. 2A).

**Figure 2:**
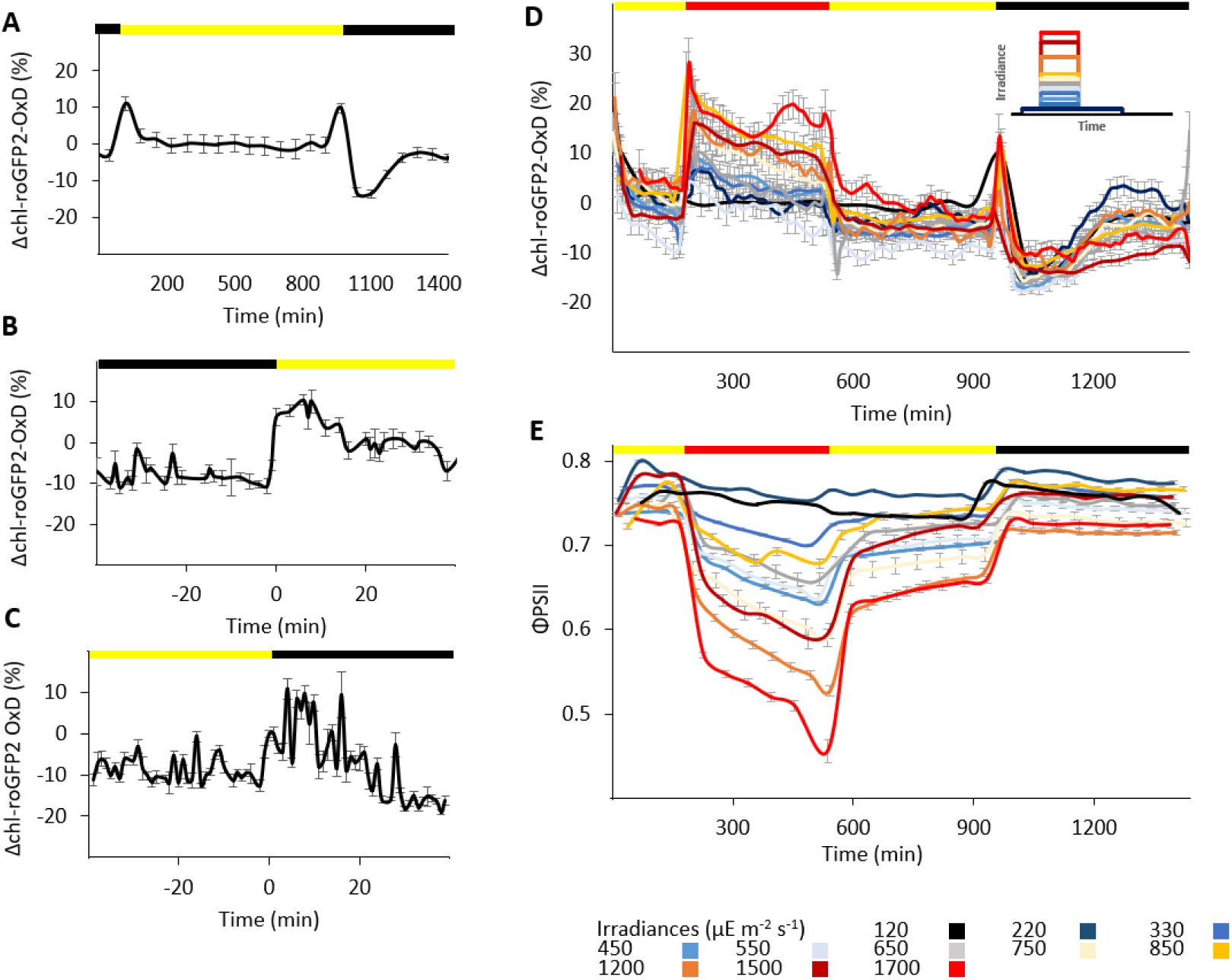
Daily changes in the degree of chl-roGFP2 oxidation (chl-roGFP2 OxD) under normal and high-light growth conditions. chl-roGFP2 fluorescence was monitored over a 24-hour period and chl-roGFP2 OxD values were calculated. Oxidation values were normalized to steady state values (as observed during the day at constant light conditions of 120µE m^-2^ s^-1^). (A) Changes in chl-roGFP2 OxD as a function of the time from the experiment onset (08:30), under normal light conditions. Values are presented as means (of 8 plants) ± SE (B) Oxidation and re-reduction of chl-roGFP2 during darkness-light transitions. Values are shown as means (of 5-24 plants, taken from 7 experiments) ± SE. (C) Oxidation and re-reduction of chl-roGFP2 during light-darkness transitions, shown as means (of 6-45 plants, taken from 20 experiments) ± SE. (D) Daily changes in chl-roGFP2 OxD during high-light experiments in wild type (WT) plants. chl-roGFP2 *Arabidopsis* plants were subjected to the following light conditions during a 24-hour period: 3 hours 120µE m^-2^ s^-1^, 6 hours 220,330,450,550,650,750,850,1200,1500 or 1700µE m^-2^ s^-1^, 7 hours 120µE m^-2^ s^-1^ and 8 hours 0µE m^-2^ s^-1^. For each treatment, between 21 and 24 plants, divided into 3 independent plates (for the 550µE m^-2^ s^-1^ experiment 2 plates were used with 7-8 plants each), were tested. For clear presentation of the oxidation trends along the x-axis, a “sliding window” approach was taken, in which each data point represents the average of three plates (n=3). Values represent means ± SE. Inset: Experimental design for the high-light experiments. (E) Daily changes in chl-roGFP2 ΦPSII values in plants exposed to the same conditions as in (D). ΦPSII values were derived from chlorophyll fluorescence analysis and represent means (of 12 plants) ± SE.

To further characterize the oxidation occurring during darkness-light transitions, we collected data from 20 experiments, cumulatively involving approximately 2600 plants. This large dataset allowed for high-resolution analysis of the chl-roGFP2 OxD values that surround the transitions’ points. We observed a rapid burst (less than 1 min) in chl-roGFP2 OxD levels, which remained stable for 10-15 min during the transition to light, and was followed by chl-roGFP2 reduction, stabilizing at higher oxidation levels than those detected during the dark period (Fig. 2B). Contrastingly, chl-roGFP2 oxidation occurred approximately 5 min after the transition from light to dark, and was followed by a gradual reduction, reaching values that were relatively lower than those detected during the day (Fig. 2C). These results demonstrate significant chl-E_GSH_ oxidation during darkness-light transitions and imply the transmission of oxidative signals during the induction and termination phases of photosynthesis.

Next, we investigated chl-roGFP2 OxD patterns under HL conditions by exposing plants to various HL intensities during a 24h cycle. Plants were exposed to three hours of normal growth light conditions (120µE m^-2^ s^-1^) at the beginning of the day, after which, the light irradiance was raised to various intensities (220, 330, 450, 550, 650, 750, 850, 1200, 1500 and 1700µE m^-2^ s^-1^) for a 6h period. After the HL phase, the light was returned to the normal intensity for an additional 7h period, followed by an 8h darkness period. (Fig. 2D inset). In addition to chl-roGFP2 oxidation (Fig. 2D), the PSII operating efficiency (ΦPSII) was continuously monitored based on chlorophyll fluorescence imaging (Fig. 2E). Immediate chl-roGFP2 oxidation, followed by gradual reduction, was observed when plants were shifted from 120 µE m^-2^ s^-1^ to any higher light intensity, indicating a rapid chl-E_GSH_ response to unpredicted increases in the light input (Fig. 2D). Reduction at the onset of darkness, followed by gradual oxidation, as observed during normal growth conditions, was observed for all light treatments (Fig. 2D).

Examination of the correlation between chl-roGFP2 oxidation and light irradiance, revealed a binary response rather than a quantitative correlation; irradiance values of 220-650µE m^-2^ s^-1^ triggered a similar initial chl-roGFP2 ΔOxD of approximately 10%, while for irradiances of 750-1700µE m^-2^ s^-1^, the initial oxidation was 20-30%. In contrast, the decrease in ΦPSII correlated with the light intensity (Fig. 2E). This binary response suggests a light-dependent threshold of chl-roGFP2 oxidation around 750μE m^−2^ s^−1^, which matched the light saturation area, as indicated by light-response curves generated from chlorophyll fluorescence and carbon assimilation measurements (Supp. Fig. 2). Overall, the data indicate the differential response of photosynthesis efficiency and redox metabolism to HL and imply a regulatory role of chl-E_GSH_ during the shift from light-unsaturated to light-saturated photosynthesis.

### Patterns of chloroplastic E_GSH_ under fluctuating light conditions

As plants are naturally exposed to discontinuous solar energy, we further examined chl-roGFP2 dynamics under fluctuating light (FL) conditions (Fig. 3). To this end, plants were exposed, in the middle of the day, to 6 hours of FL between 1700µE m^-2^ s^-1^ and 120µE m^-2^ s^-1^, at frequencies of 1, 5 and 10 min (Fig. 3A, D and G). Under all tested frequencies, a gradual and partial loss of photosynthetic activity was observed, as indicated by decreasing ΦPSII values, reaching a minimum at the end of the FL period (Supp. Fig. 3). Shifting plants back to normal growth light conditions resulted in partial ΦPSII recovery, with values still lower than those measured at the beginning of the day (Supp. Fig. 3). Plants exposed to the 1 min frequency showed a pattern that resembled HL treatments, including an initial chl-roGFP2 (ΔOxD = ∼20%, Fig. 3B), followed by a gradual reduction when plants were moved back to normal growth light conditions. In contrast, oscillations in the chl-roGFP2 oxidation were observed in plants exposed to frequencies of 5 min and 10 min (Fig. 3E and H).

**Figure 3:**
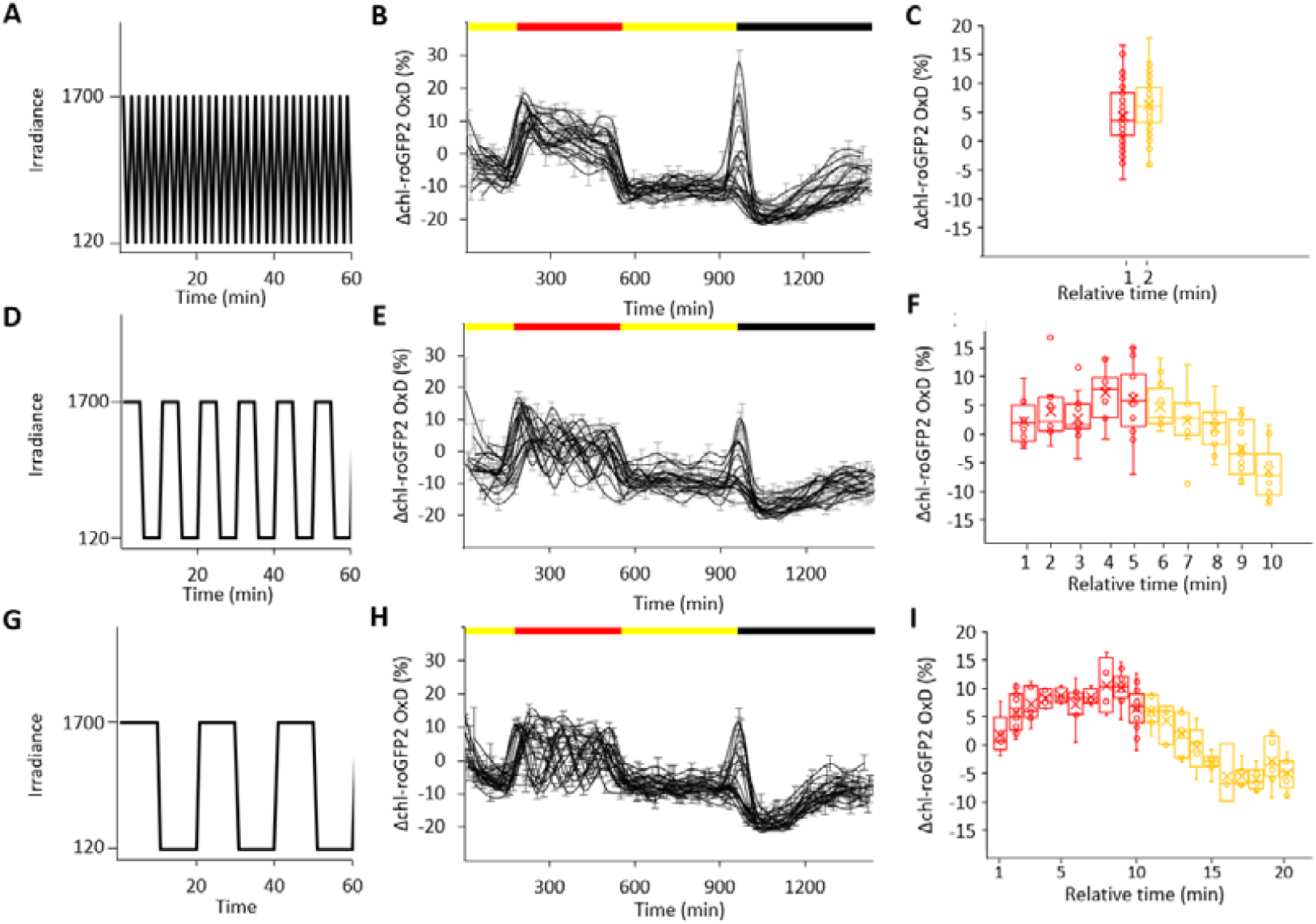
Daily changes in chl-roGFP2 OxD under fluctuating light conditions. *Arabidopsis* plants were subjected to the following light conditions during a 24-hour period: 3 hours 120µE m^-2^ s^-1^, 6 hours “fluctuating light”, 7 hours 120µE m^-2^ s^-1^ and 8 hours 0µE m^-2^ s^-1^. Fluctuating light ranged between 120-1700µE m^-2^ s^-1^, delivered at a frequency of 1, 5 or 10 min. The light irradiance as a function of the time, over a 60-minute period, extracted from the fluctuating light phase is presented for the applied frequencies of 1, 5 and 10 min (A, D and G, respectively). Changes in chl-roGFP2 OxD values as a function of the time (min) from the experiment onset (08:30) (B, E and H). Oxidation values were normalized to steady state values (as observed during the day at constant light conditions of 120µE m^-2^ s^-1^). Values represent means (of 8 plants) ± SE. ΔOxD as a function of relative time (min) within each fluctuating light cycle (C, F and I). The values taken from the 1700µE m^-2^ s^-1^ period within each cycle are in presented the red box plots, while the values taken from the 120µE m^-2^ s^-1^ within each cycle are presented in yellow box plots, both representing means of 6-101 plants ± SE.

To simplify the visual analysis, all the values recorded during the FL treatments were combined into a cycle-like presentation, *i.e.*, all minutes are shown as relative to their cycle (*e.g.* under the 10 min frequency, both 8:00 and 8:20 were considered the 20^th^ minute, each for their own cycle, and values observed in both of them were pooled together). A cyclical oscillation between higher and lower chl-roGFP2 oxidation states was noted during the cycles of 5 min and 10 min frequencies, but not for the 1 min frequency cycles, with the latter showing no significant difference between values recorded under the HL phase and those recorded under the ‘normal-light’ phase (equivalence test at Δ=5, p <0.05). Importantly, these patterns were conserved in all FL cycles, and were not altered, despite the gradual decrease in ΦPSII during the FL phase (Supp. Fig. 3). Taken together, these results point to frequency-dependent relaxation of the chl-E_GSH_ under FL.

### Chloroplastic E_GSH_ dynamics in mutants impaired in photoprotective mechanisms

To examine the interaction between chl-E_GSH_ patterns and light photoprotection mechanisms, the chl-roGFP2 probe was expressed in *npq1* and *pgr5* plants and chl-roGFP2 OxD was measured in parallel with ΦPSII under the same HL regimens (Fig.4). Despite the clear decrease in ΦPSII in *npq1*, as compared to WT plants under identical HL conditions (*e.g.*, approximately 0.3 and 0.5 under 1700 µE m^-2^ s^-1^, respectively; Fig. 4D, E), nearly identical chl-roGFP2 oxidation trends were observed in both lines (Fig. 4A, B). Highly impaired photosynthetic efficiency was observed in *pgr5* plants exposed to HL, reaching ΦPSII values of approximately 0.15 (Fig. 4F). As in WT, shifting of *pgr5* to HL resulted in fast oxidation, but chl-roGFP2 OxD values recovered much quicker than in WT (Fig. 4C and Fig. 4A, respectively). These results suggest that photoinhibition of PSII does not necessarily result in higher production of ROS and that PSI photoinhibition results in decreased ROS production.

**Figure 4:**
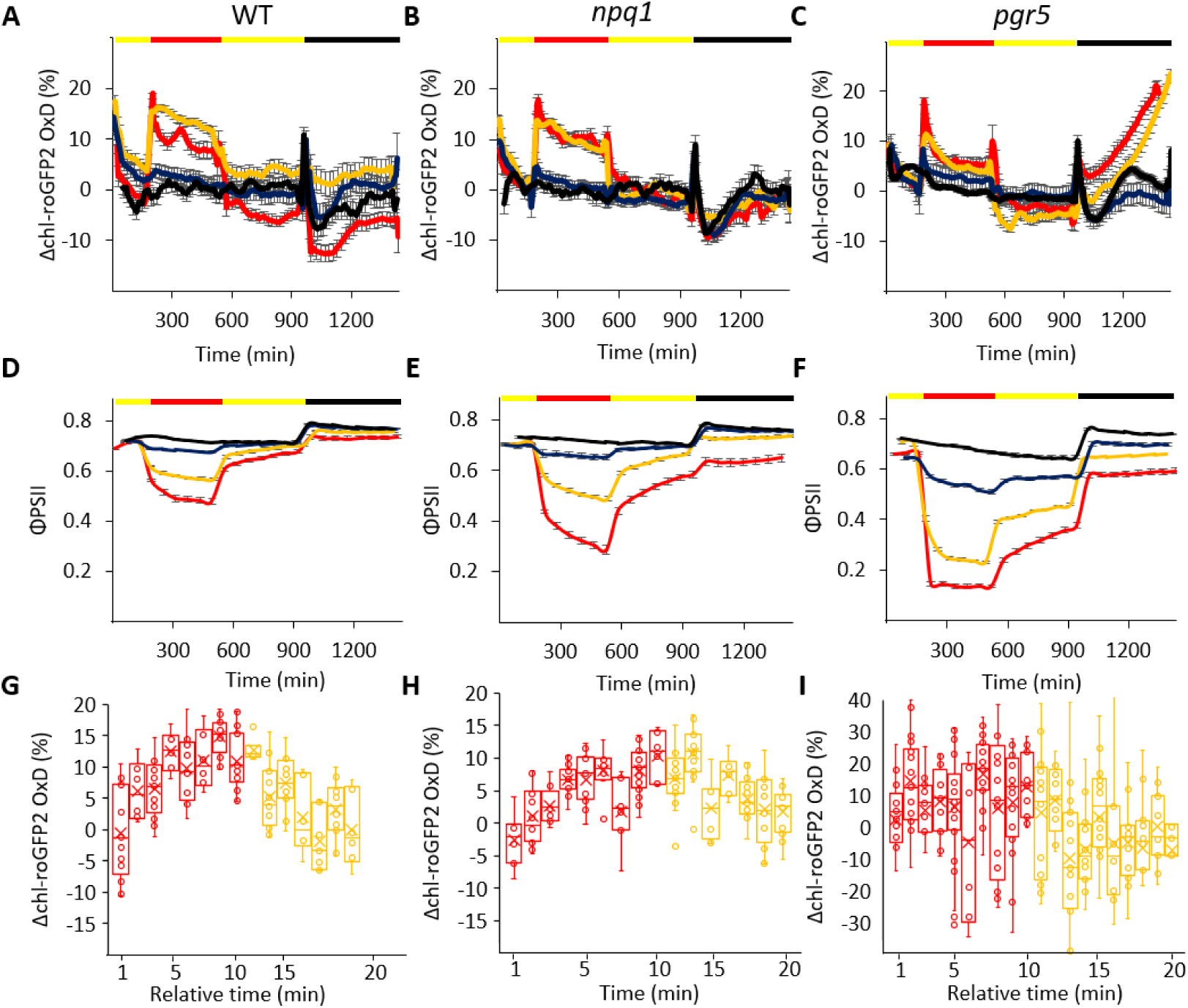
Daily changes in chl-roGFP2 OxD and ΦPSII under high-light and fluctuating light conditions in photoprotective mutants. WT, *npq1* and *pgr5 Arabidopsis* plants were subjected to the following high-light conditions during a 24-hour period: 3 hours 120µE m^-2^ s^-1^, 6 hours 120, 220, 850 or 1700µE m^-2^ s^-1^ (black, blue, yellow and red lines, respectively, in A-F), 7 hours 120µE m^-2^ s^-1^ and 8 hours 0µE m^-2^ s^-1^. chl-roGFP2 fluorescence was monitored throughout the experiment and chl-roGFP2 OxD values were calculated. Changes in chl-roGFP2 OxD as a function of the time from the experiment onset is presented for WT (A), *npq1* (B) and *pgr5* (C) plants. Oxidation values were normalized to steady state oxidation levels. Each treatment involved between 24 and 32 plants divided into four independent plates that were consolidated in a “sliding window” (n=4) display. Values represent means ± SE. ΦPSII as a function of the time from the experiment onset is presented for WT (D), *npq1* (E) and *pgr5* (F) plants. ΦPSII values were derived from chlorophyll fluorescence analysis and represent means (of 12 plants) ± SE. (G-I) *Arabidopsis* plants were subjected to the fluctuating light experiments as described in Fig. 3G. Changes in chl-roGFP2 OxD as a function of relative time (min) within each 10-minute frequency fluctuating light cycle is presented for WT (G), *npq1* (H) and *pgr5* (I) plants. The values taken from the 1700µE m^-2^ s^-1^ period within each cycle are presented in red box plots, while the values taken from the 120µE m^-2^ s^-1^ within each cycle are presented in yellow box plots, both representing means of 6-26 plants± SE.

As PGR5 was found to be essential for photoprotection, specifically under FL conditions (Munekage et al., 2002; Suorsa et al., 2013; Yamori et al., 2016; Yamamoto and Shikanai, 2019), we examined chl-roGFP2 oxidation in WT, *npq1* and *pgr5* plants under FL of frequencies of 1 min and 10 min, as explained earlier. The observed whole-day oxidation patterns are depicted in Supp. Fig. 4 and the cyclic presentations are shown in Fig. 4. Under the 1 min frequency regimen, no significant difference in oxidation patterns was observed between the HL phase and the normal-light phases of each cycle in all three lines (using the equivalence test at Δ=5%, *p*-value=0.05; Supp. Fig. 5). Contrastingly, under the 10-min frequency regimen, the chl-roGFP2 redox state oscillated between relatively reduced and relatively oxidized values, according to the relative timing within the FL cycle. Intriguingly, this phenomenon occurred only in WT and *npq1* plants and not in *pgr5* plants (Fig. 4G-I; the ΦPSII data is presented in Supp. Fig. 6). Taken together, these results demonstrate that the chl-E_GSH_ response to FL conditions is dependent on PGR5 activity and is impaired under PSI photodamage.

### Gradual relaxation of chloroplastic E_GSH_ oxidation and ΦPSII impairment in *pgr5* plants

Interestingly, a significant increase in chl-roGFP2 OxD was observed in *pgr5* plants during the night following high and FL conditions. As shown in Figure 4C (and Supp. Fig. 4), oxidation levels gradually increased during the night, reaching levels that correlated with the light intensity applied the day before. Accordingly, higher oxidation values were observed during the night following HL treatments of 1700 m^-2^ s^-^ compared to 850µE m^-2^ s^-1^ (Figure 5). After the night, chl-roGFP2 OxD values returned to steady state OxD levels, immediately when light was turned on.

**Figure 5:**
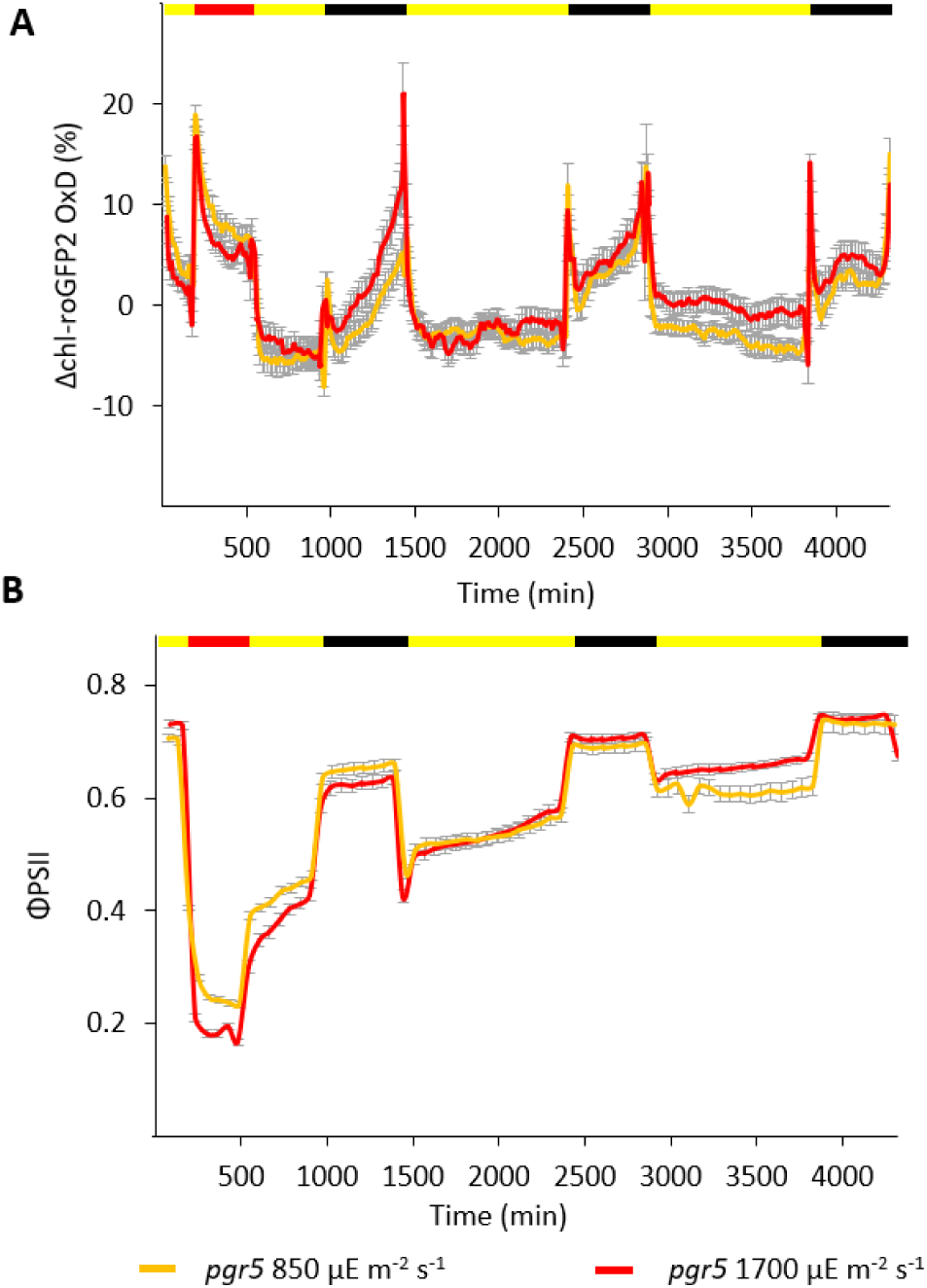
Daily changes in chl-roGFP2 OxD and ΦPSII during one day of high-light regime followed by two days of normal light conditions in *pgr5* plants. *Arabidopsis pgr5* plants were subjected to the following light conditions during a 72-hour period: 3 hours 120µE m^-2^ s^-1^, 6 hours 850 or1700µE m^-2^ s^-1^, 7 hours 120µE m^-2^ s^-1^, 8 hours 0µE m^-2^ s^-1^, followed by two 24-hour cycles of: 16 hours 120µE m^-2^ s^-1^ and 8 hours 0µE m^-2^ s^-1^. (**A**) Changes in chl-roGFP2 OxD as a function of the time from the experiment onset. chl-roGFP2 fluorescence was monitored throughout the experiment and chl-roGFP2 OxD values were calculated. Oxidation values were normalized to steady state oxidation levels. Each treatment involved 32 plants divided to four independent plates that were consolidated in a “sliding window” (n=4) display. Values represent means (of 8-32 plants) ± SE. (**B**) ΦPSII as a function of time from the experiment onset. ΦPSII values were derived from chlorophyll fluorescence analysis and represent means (of 12 plants) ± SE. Data of similar experiment performed with *gl1* lines, which served as the reference ecotype for the *pgr5* line, are depicted in Supp. Fig. 7.

To further elucidate this phenomenon, we subjected *pgr5* plants to 72h experiments consisting of 24h HL experiments (containing a 6h HL period of 850 and 1700 µE m^-2^ s^-1^), followed by two normal-light days (Fig. 5A, B; *gl1* data, which served as the background line for the *pgr5* mutation, are presented in Supp. Fig. 7). During the first day of recovery under normal-light conditions, ΦPSII values were still lower than in WT plants (0.5-0.56 versus approximately 0.7), and night chl-roGFP2 OxD oxidation clearly occurred again, albeit to a lower degree. Complete relaxation of the night chl-roGFP2 OxD oxidation was achieved only during the third night after HL treatment, reaching chl-roGFP2 OxD and ΦPSII values comparable to those observed under normal-light conditions (Supp. Fig. 8). During the days, despite night oxidation, the chl-roGFP2 values were comparable on all the examined days. This night oxidation effect, which appeared hours after the HL stress, suggests a link between photoinhibition during the day and alterations in redox metabolism during the nights following the stress conditions.

## Discussion

Light-dependent flow of electrons from water to ferredoxin, the photosynthetic linear electron flow (LEF), produces the reducing power required for diverse downstream metabolic events, including carbon and nitrogen assimilation, as well as for antioxidant activity. On the other hand, reduction of molecular oxygen at the acceptor site of PSI is the main process of superoxide anion radical (O_2_^.−^) and H_2_O_2_ production in photosynthesizing chloroplasts (Asada, 1999; Osmond et al., 2000). Thus, solar energy is the source for both reducing power and oxidizing agents, subsequently requiring tight regulation to avoid imbalances between the two.

Monitoring photosynthesis-derived H_2_O_2_ production is challenging, with current approaches (for the estimation of the WWC activity flux) based mainly on uptake measurements of isotopically labeled O_2_ (Driever and Baker, 2011; Gauthier et al., 2018; Holloway-Phillips, 2018). Despite the clear occurrence of the Mehler reaction (Mehler, 1951; Asada, 1999), only few studies have actually managed to measure O_2_ photoreduction *in vivo*, thus hindering detailed investigation of its physiological role. Owing to its high sensitivity and reversibility, the recently developed genetically-encoded, redox-sensitive roGFP probes allow *in vivo* mapping of the dynamics of redox metabolism in subcellular compartments. The ascorbate-glutathione pathway and glutathione peroxidase activity are the main factors influencing H_2_O_2_ reduction and E_GSH_ redox state (Rahantaniaina et al., 2013). Accordingly, measurements of light-dependent alterations in chl-E_GSH_ can reflects the balance between photosynthesis-dependent H_2_O_2_ production and antioxidant activity. Though probe oxidation can also be mediated directly by H_2_O_2_, these two scenarios are physiologically similar, as roGFP mimics natural chloroplastic redox-sensitive proteins, which receive reduction and oxidation signals from the photosynthetic machinery (Nietzel et al., 2019).

By measuring chloroplast-targeted roGFP2 oxidation at high temporal resolution, over several consecutive days and in response to diverse light conditions, this work uncovered several patterns of photosynthesis-dependent ROS production. The presented results showed oxidation peaks during darkness-light and light-darkness transitions (Fig. 2B, C). During the former, the discrepancy between the light-dependent reactions, which are activated immediately upon light exposure, and between the Calvin cycle reactions, which involves a 15 min activation phase, resulted in limited overflow of electrons from PSI to NADP^+^ (Wirtz et al., 1982; Tikhonov, 2015) and transient diversion of electrons to O_2_ reduction. The subsequent relaxation of the chl-roGFP2 oxidation state could have resulted from the full induction of CO_2_ assimilation activity and decrease in O_2_ reduction in parallel with H_2_O_2_ detoxification via antioxidants. These results are in line with the suggested induction of the WWC during the induction phase of photosynthesis (Radmer and Kok, 1976; Miyake, 2010; Volpert et al., 2018).

The light-dependent oxidants, which trigger changes in chl-E_GSH_, may transmit oxidative signals to redox-sensitive proteins during the photosynthesis induction phase (Buchanan and Balmer, 2005; Rahantaniaina et al., 2013). Indeed, reception of oxidative signals by chloroplastic atypical thioredoxins (which regulate NPQ induction and starch biosynthesis) was demonstrated shortly after illumination (Dangoor et al., 2012; Eliyahu et al., 2015). Similarly, peaks in oxidant production during light-darkness transitions can play a regulatory role in oxidizing redox-sensitive proteins at night, allowing transmission of reducing signals when the light is turned on again. Oxidative signals transmitted from H_2_O_2_ to target proteins, mediated by 2-Cys peroxiredoxin at the onset of the dark period, have been demonstrated (Eliyahu et al., 2015; Ojeda et al., 2018; Vaseghi et al., 2018). Similarly, protein oxidation of key proteins in photosynthesis, including ATP synthase CF1-γ subunit, fructose 1,6-bisphosphatase (FBPase) and sedoheptulose 1,7-bisphosphatase (SBPase), has been observed after the light was switched off (Yoshida et al., 2014). However, the mechanism underlying the induction of these oxidation peaks during light-darkness transitions, is not clear.

The WWC has been suggested to play a role in the dissipation of excess photon energy, thereby protecting PSI from photoinhibition under environmental stresses (Biehler and Fock, 1996; Asada, 1999; Ort and Baker, 2002; Miyake, 2010). Although the high proportion of photochemically absorbed electrons resulting in O_2_ reduction in cyanobacteria and diatoms (up to 50%) can support this physiological role, the limited capacity of WWC in C3 plants indicates that it does not act as a dissipation mechanism therein (Osmond et al., 2000; Waring et al., 2010). Therefore, it has been suggested, that in higher plants, it plays a physiological role in transmitting light□induced redox signals (Driever and Baker, 2011; Gerken et al., 2020). Indeed, Exposito-Rodriguez *et al.* (Exposito-Rodriguez et al., 2017) demonstrated, using the H_2_O_2_-sensitive HyPer2 probe, that photosynthesis-dependent H_2_O_2_ can influence the nuclear redox state and gene transcription, providing a HL acclimation mechanism.

Our data showed that, as opposed to a relatively reduced ‘steady state’ under normal growth conditions, chl-roGFP2 underwent considerable oxidation under HL conditions (Fig. 2D). Interestingly, light irradiances of 220-650µE m^-2^ s^-1^ induced a lower chl-roGFP oxidation than irradiances of 750-1700µE m^-2^ s^-1^, while within the two ranges, there did not seem to be a significant difference. This binary effect, together with the contrasted non-binary behavior of the drop in ΦPSII during the HL period, suggests a regulatory role of photosynthesis–derived oxidation, rather than a safety valve for energy dissipation. Furthermore, it also implies an existence of a “threshold” around the light saturation point (Supp. Fig. 2), which separates between two distinct states of oxidant production and, consequently, the chl-E_GSH_ redox state. We hypothesize that these two redox states signal distinct metabolic responses, in accordance with the severity of the HL conditions. The similarity in chl-E_GSH_ OxD patterns between WT and *npq1* plants, despite the significantly lower PSII efficiency in the mutant line (Fig. 4), further supports this signaling role of ROS production, as non-photochemical quenching is a major mechanism for dissipating excess photons in higher plants (Osmond et al., 2000).

A frequency-dependent shift between oxidation states was observed under FL conditions (Fig. 3), suggesting its role in photosynthesis regulation under natural conditions, in which the solar flux varies due to the diurnal cycle, canopy structure and varying cloud coverage. Interestingly, while a decrease in photosynthesis efficiency was observed during FL (Supp. Fig. 3), the ability to shift between the two chl-roGFP2 oxidation states was not impaired throughout the FL period (Fig. 3). The fact that the observed fluctuating oxidation patterns were not induced in *pgr5* plants points towards the connection between PSI photoinhibition and rates of ROS production under these conditions. This was also observed under HL conditions, whereas faster reduction was observed in *pgr5* plants (Fig. 4C). Recently, it has been demonstrated that *pgr5* plants are highly sensitive to HL-induced damage of iron– sulphur (Fe-S) clusters of PSI and that this damage provides an additional photoprotective mechanism by inducing a non-photochemical photoprotective energy quenching state (Tiwari et al., 2016). Based on the data observed here, we suggest that this PSI quenching state results in decreased oxygen reduction via the WWC cycle. This results are in line with the suggested role of PSI photoinhibition in preventing excessive ROS production (Lima-Melo et al., 2019a) and the increased antioxidant capacity and decreased production superoxide and H_2_O_2_ in *pgr5* plants (Suorsa et al., 2012). Accordingly, it is possible that alterations in ROS production and downstream redox signaling affects the sensitivity of *pgr5* to FL conditions.

While focusing on light-dependent oxidant production, continuous diurnal chl-roGFP2 OxD measurement uncovered a unique phenotype in *pgr5* plants during the nights following the HL and FL conditions. Severe gradual oxidation was observed in the first night that followed the HL stress, reaching maximum oxidation by the end of the night (Fig. 4C). Furthermore, night-time oxidation was observed again the next night and typical night redox patterns seen in WT plants were only observed in *pgr5* plants on the third night after treatment (Fig. 5A). These effects, which only surfaced hours after exposure to light stress, were not directly caused by photoinhibition and may be related to the slow recovery time of PSI (Sonoike, 2011; Zhang et al., 2011). Accordingly, restoration of CO_2_ assimilation in HL-treated *pgr5* plants was achieved during the third day in normal light conditions and was attributed to recovery of PSI parameters, such as maximum oxidizable P700 and maximal reduction state of ferredoxin (Lima-Melo et al., 2019b).

It is possible, that chl-E_GSH_ oxidation, observed during the two nights subsequent to HL stress, is a result of starch starvation caused by PSI photoinhibition in *pgr5* plants (Lima-Melo et al., 2019b). It has been reported that starch accumulates in chloroplasts during the day and is metabolized during the night as a source of NADPH, via the pentose phosphate pathway (Weise et al., 2004; Kirchsteiger et al., 2009). Therefore, it is possible that the observed night-time oxidation in *pgr5* plants following HL stress is due to a deficiency in NADPH as a result of starch deficiency. However, the demand for NADPH to maintain the highly reduced state of chl-E_GSH_ during the night, assumes GSH oxidation at night, which, to our knowledge, has not yet been reported. It is possible that the oxidation at night may be due to export of GSH from the chloroplasts to the mitochondria, which are the main source of ROS during the night, resulting in a lower GSH/GSSG ratio in chloroplasts

Interestingly, recent works demonstrated that in addition to their role in thiol-redox maintenance, GSH and glutaredoxins join to assemble Fe–S clusters and transfer them to acceptor proteins, linking between GSH and iron metabolism (Rouhier et al., 2008; Mühlenhoff et al., 2010; Rouhier et al., 2010; Kumar et al., 2011). Accordingly, repair and biogenesis of PSI after HL-induced damage of Fe-S clusters may require the function of GSH and glutaredoxins. It is possible that the observed oxidation of chl-E_GSH_ several nights after HL or FL stress in *pgr5* plants (Fig. 4C, Fig. 5 and Supp. Fig.4), in which PSI centers are particularly sensitive to photoinhibition, mirrors the divert of reduced GSH for the slow process of biogenesis and repair of PSI Fe-S clusters. In view of that, the gradual oxidation of chl-E_GSH_ at night, as observed during normal growth conditions as well as during HL and FL conditions in WT plants (Fig. 2D and Supp. Fig. 4), reflects the natural repair daily cycle of PSI. Whether this repair machinery operates exclusively at nights, or that the shortage of photosynthetically production of reducing power in the dark allowed to uncover this GSH requirement is not clear. A conflict between these two functions of GSH may occur under chilling stress conditions, in which PSI is the major site of photoinhibition (Terashima et al., 1994), resulting in slow repair of PSI Fe-S clusters that will be further aggravated when glutathione reductase activity is suppressed (Shu et al., 2011). Further work will be required to characterize the molecular mechanisms regulating nighttime chl-E_GSH_ oxidation and the possible connection to PSI recovery.

In conclusion, by continuously monitoring chloroplast-specific E_GSH_ oxidation patterns over several days, this work provides a comprehensive view on photosynthetically produced ROS under various light conditions and uncover an uncharacterized link between PSI photoinhibition and night GSH metabolism. High-resolution monitoring of compartment-specific redox metabolism under varying environmental conditions will further expand the fundamental understanding of the role of redox signaling in light acclimation of higher plants.

## Material and Methods

### Plant Material, Growth Conditions and Experimental Set-up

*Arabidopsis thaliana* **WT** (ecotype Columbia-0), ***npq1*** (CS3771, At1g08550, obtained from ABRC), and ***pgr5*** (EMS mutant line, At2g05620, obtained from Prof. T. Shikanai; lines were used throughout this research. Columbia ecotype *glabrous 1* (gl-1; obtained from Prof. T. Shikanai [Kyoto University]), which served as the reference ecotype for the *pgr5* line, was also used. Plants were sown on soil, put in 4°C for two days and grown under 16/8 light-dark cycles with photosynthetic photon flux density (PPFD) of 120µE m^-2^ s^-1^ (21°C, 60-70% RH, ambient CO_2_) for two-three weeks. For roGFP and PAM analysis, plants were transferred to 12-Well Cell Culture Multiwell Plates in solid peat plugs and the plugs were covered (with holes for plants) with black plastic to prevent autofluorescence. For 3-(3,4-dichlorophenyl)-1,1-dimethylurea (DCMU) experiments, 2-3-week-old plants were treated with 150µM DCMU (D2425-100G, Sigma), kept for 1 hour in the dark and then exposed to high light (HL) conditions of 700µE m^-2^ s^-1^.

In all experiments, 2-3-week-old plants were incubated in 21°C, 60-70% RH and ambient CO_2_. The following light conditions were applied: Normal growth experiments were under 16 hours of 120µE m^-2^ s^-1^ / 8 hours of 0µE m^-2^ s^-1^ (24-hour cycle). For HL experiments, plants were exposed to 3 hours of normal light conditions (120µE m^-2^ s^-1^), and then light irradiance was elevated during 6 hours to the following HL intensities: 220, 330, 450, 550, 650, 750, 850, 1200, 1500 or 1700µE m^-2^ s^-1^. After the high-light phase, plants were grown again under normal light conditions for an additional 7 hours and finally were under an 8-hour darkness period. For fluctuating light (FL) experiments, plants were exposed to 3 hours of 120µE m^-2^ s^-1^, followed by 6 hours of fluctuating light between 1700 (HL) and 120 (normal light) µE m^-2^ s^-1^ in frequencies of 1,5 or 10 minutes (*i.e.* 1,5 or 10 minutes of 1700µE m^-2^ s^-1^ followed by 1,5 or10 minutes 120µE m^-2^ s^-1^ and vice versa *etc.*). Afterwards, plants were grown again for 7 hours under 120µE m^-2^ s^-1^ and finally under an 8-hour darkness period.

### Generation of chl-roGFP2 expressing lines

Chloroplast targeting was achieved by using either the Transketolase signal peptide (33) or a 2-Cys Peroxiredoxin A signal peptide (this study), both target proteins to the chloroplast stroma (König et al., 2002; Schwarzländer et al., 2008). For the generation of the chl-roGFP2 line, the full gene sequence was chemically synthesized. The first 74 amino acids from chloroplastic Peroxiredoxin A (PRXa; Uniprot ID: Q96291) were used as a signal peptide. The chl-roGFP2 gene was cloned into the plant cloning vector pART7 using *Xho*I and *Hin*dIII restriction enzymes. The whole construct including the CaMV 35S promoter and ocs terminator was then cloned into the binary vector pART27 using the restriction enzyme *Not*I. The pART27 plasmid, which contains the chl-roGFP2 construct, was transformed into GV3101 *Agrobacterium tumefaciens.* Transformation of *Arabidopsis thaliana* (Columbia, Col-0) was performed by floral dip (Clough and Bent, 1998). Transformants lines were selected based on Kanamycin resistance and the chl-roGFP2 fluorescence signal.

### Confocal microscopy

Images were acquired with a Leica TCS SP8 confocal system (Leica Microsystems) and the LAS X Life Science Software, while using a HC PL APO ×40/1.10 objective. All images were acquired with at a 4096 × 4096-pixel resolution. Images were acquired as the emission at 500-520nm following excitation at 488nm for chl-roGFP2 fluorescence and emission at 670nm following excitation at 488nm for chlorophyll fluorescence. Merged images were generated using Fiji (Image J) software.

### chl-roGFP2 Fluorescence Measurements and Analysis

Whole-plant chl-roGFP2 fluorescence imaging was detected using an Advanced Molecular Imager HT (Spectral Ami-HT, Spectral Instruments Imaging, LLC., USA), and AMIview software for image acquisition. This analysis was used to screen for fluorescent plants and for qualitative analysis. For chl-roGFP2 fluorescence detection, excitation using 405nm/20 or 465nm/20 LED light sources and 510nm/20 emission filter were used. For chlorophyll detection, 405nm/20 LED light source and 670nm/20 emission filter used were. Quantitative chl-roGFP2 analysis was mainly carried out by the fluorometer method (Rosenwasser et al., 2010) using a Tecan Spark® multimode microplate reader. For all results, the following filters were used: 400nm/20 (excitation) / 520nm/10 (emission) and 485nm/20 (excitation) / 520nm/10 (emission). For chlorophyll detection, the following filters were used: 400nm/20 (excitation) / 670nm/40 (emission). For automatic detection of chl-roGFP2 signals, plants growing in 12-well plates were placed in a PSI Fytoscope FS-SI-4600 chamber and were automatically inserted into the fluorometer for fluorescence detection, using a paa automations KiNEDx KX.01467 robot, throughout 24-hour, 48-hour and 72-hour experiments using a self-compiled paa automations Overlord™ program.

For the plate-reader analysis, from each well a 9-by-9-pixel matrix was formed. chlorophyll fluorescence was detected, in order to form a chlorophyll mask, which was used to choose pixels which returned a positive chlorophyll fluorescence signal, and only those pixels were taken into account for the roGFP analysis. Then, the averages of non-fluorescent plants (WT) was calculated, and the values were used to be subtracted as background autofluorescence from the values detected in the chl-roGFP2 fluorescence analysis. roGFP2 degree of oxidation (the relative quantity of oxidized roGFP proteins, OxD) was calculated based on the fluorescence signal according to Equation 1 (Meyer et al., 2007):

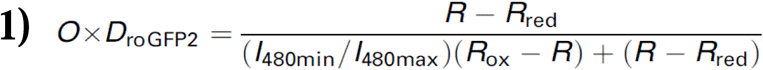

where R represents the ratio (405/488) at each point in the experiment, R_red_ represents the ratio under fully reduced conditions, R_ox_ represents the ratio under fully oxidized conditions, I480_min_ (or I488_ox_) represents the fluorescence emitted at 510nm when excited at 488nm under fully oxidized conditions and I480_max_ (or I488_red_) represents the fluorescence emitted at 510nm when excited at 488nm under fully reduced conditions. In order to be able to calculate chl-roGFP2 OxD, we subjected chl-roGFP2 *Arabidopsis* plants and measured the 400/480nm fluorescence ratio under fully oxidized and fully reduced conditions, by using 1-4M H_2_O_2_ and 100mM DTT, respectively. A self-compiled Matlab script was written to allow robust analysis of many files which contain the measurements in each experiment. The scripts collect the roGFP and chlorophyll fluorescence values from each plant into one matrix. In addition, experimental metadata such as the beginning and ending time, plant lines, light irradiance, temperature, RH (%) and CO_2_ concentration were also collected and added to the output file.

### Chlorophyll Fluorescence and Gas Exchange Measurements

ΦPSII of each line was detected using a Walz PAM IMAGING PAM M-*series* IMAG-K7 (MAXI) Fluorometer (a plate of 12 plants was measured every 60 minutes). ETR was calculated as the multiplication of ΦPSII by the quantity of photons emitted by the light source (PAR), by the average percentage of the photons absorbed by the photosynthetic machinery (0.84), by the amount of photons allocated to PSII (0.5; assuming that PSI and PSII contain the same amount of reaction centers, therefore each photosystem receives 50% of the photon influx). Accordingly, ETR= Y(II) * PAR * 0.84 * 0.5 (Maxwell and Johnson, 2000). Electron transport rate (ETR) was measured and calculated using the Walz PAM Fluorometer. A gas exchange light induction curve (Carbon Assimilation as a function of light irradiance) was performed using the LI-6800 (LI-COR^®^). 5-6 plants were planted in LI-COR^®^ 65mm pots (610-09646) and analysis was done using the LI-6800 small plant chamber (6800-17). Total leaf area was calculated via image analysis using the Ami HT Imager and Matlab. Full ETR and Carbon assimilation light induction curves’ data are in Supp. Fig. 9-10.

### Statistics

Values in graphs mainly represent means of 8 plants for chl-roGFP2 OxD and 12 plants for ΦPSII, and error bars represent the respective standard errors. For many experiments, 3-4 plates were analyzed and for a clear presentation, a “sliding window” approach was taken, in which each data point represents the average of three-four plates. Statistical significance was mainly tested using a two-tail Student’s t-test, equivalence test significance and analysis of variance (ANOVA) at 95% Confidence Level and was indicated by asterisks when shown.

## Author contributions

Z.H., and S.R. designed research, performed research, analyzed data and wrote the manuscript. S.R supervised the study.

## Acknowledgments

We thank Avihai Danon, Zach Adam, Assaf Vardi and Nardy Lampl for critical comments on the manuscript. We thank Einat Zelinger and Daniel Waiger for their help in acquiring the confocal images. We thank Prof. Toshiharu Shikanai from Kyoto University, who kindly gave us *pgr5* and *gl1* seeds for this research. This research was supported by the Israel Science Foundation (grant No. 826/17 and No. 827/17) to SR.

## References

Albrecht SC, Barata AG, Großhans J, Teleman AA, Dick TP (2011) *In vivo* mapping of hydrogen peroxide and oxidized glutathione reveals chemical and regional specificity of redox homeostasis. Cell Metab 14: 819–829

Asada K (1999) The water-water cycle in chloroplasts: scavenging of active oxygens and dissipation of excess photons. Annu Rev Plant Physiol Plant Mol Biol 50: 601–639

Avenson TJ, Cruz JA, Kanazawa A, Kramer DM (2005) Regulating the proton budget of higher plant photosynthesis. Proc Natl Acad Sci U S A 102: 9709 LP – 9713

Awad J, Stotz HU, Fekete A, Krischke M, Engert C, Havaux M, Berger S, Mueller MJ (2015) 2-Cysteine peroxiredoxins and thylakoid ascorbate peroxidase create a water-water cycle that is essential to protect the photosynthetic apparatus under high light stress conditions. Plant Physiol 167: 1592 LP – 1603

Biehler K, Fock H (1996) Evidence for the contribution of the mehler-peroxidase reaction in dissipating excess electrons in drought-stressed wheat. Plant Physiol 112: 265–272

Bratt A, Rosenwasser S, Meyer A, Fluhr R (2016) Organelle redox autonomy during environmental stress. Plant Cell Environ 39: 1909–1919

Buchanan BB, Balmer Y (2005) Redox regulation: a broadening horizon. Annu Rev Plant Biol 56: 187–220

Clough SJ, Bent AF (1998) Floral dip: a simplified method for Agrobacterium-mediated transformation of *Arabidopsis thaliana*. Plant J 16: 735–743

van Creveld SG, Rosenwasser S, Schatz D, Koren I, Vardi A (2015) Early perturbation in mitochondria redox homeostasis in response to environmental stress predicts cell fate in diatoms. ISME J 9: 385–395

Dangoor I, Peled-Zehavi H, Wittenberg G, Danon A (2012) A chloroplast light-regulated oxidative sensor for moderate light intensity in *Arabidopsis*. Plant Cell 24: 1894 LP – 1906

Dooley CT, Dore TM, Hanson GT, Jackson WC, Remington SJ, Tsien RY (2004) Imaging dynamic redox changes in mammalian cells with green fluorescent protein indicators. J Biol Chem 279: 22284–22293

Driever SM, Baker NR (2011) The water–water cycle in leaves is not a major alternative electron sink for dissipation of excess excitation energy when CO_2_ assimilation is restricted. Plant Cell Environ 34: 837–846

Eliyahu E, Rog I, Inbal D, Danon A (2015) ACHT4-driven oxidation of APS1 attenuates starch synthesis under low light intensity in *Arabidopsis* plants. Proc Natl Acad Sci 112: 12876 LP – 12881

Exposito-Rodriguez M, Laissue PP, Yvon-Durocher G, Smirnoff N, Mullineaux PM (2017) Photosynthesis-dependent H_2O2_ transfer from chloroplasts to nuclei provides a high-light signalling mechanism. Nat Commun 8: 49

Foyer CH, Noctor G (2011) Ascorbate and glutathione: the heart of the redox hub. Plant Physiol 155: 2 LP – 18

Gauthier PPG, Battle MO, Griffin KL, Bender ML (2018) Measurement of gross photosynthesis, respiration in the light, and mesophyll conductance using H_2_^18^O labeling. Plant Physiol 177: 62 LP – 74

Gerken M, Kakorin S, Chibani K, Dietz K-J (2020) Computational simulation of the reactive oxygen species and redox network in the regulation of chloroplast metabolism. PLOS Comput Biol 16: e1007102

Gutscher M, Pauleau A-L, Marty L, Brach T, Wabnitz GH, Samstag Y, Meyer AJ, Dick TP (2008) Real-time imaging of the intracellular glutathione redox potential. Nat Methods 5: 553–559

Hanson GT, Aggeler R, Oglesbee D, Cannon M, Capaldi RA, Tsien RY, Remington SJ (2004) Investigating mitochondrial redox potential with redox-sensitive green fluorescent protein indicators. J Biol Chem 279: 13044–13053

Holloway-Phillips M (2018) Photosynthetic oxygen production: new method brings to light forgotten flux. Plant Physiol 177: 7 LP – 9

Jiang K, Schwarzer C, Lally E, Zhang S, Ruzin S, Machen T, Remington SJ, Feldman L (2006) Expression and characterization of a redox-sensing green fluorescent protein (reduction-oxidation-sensitive green fluorescent protein) in *Arabidopsis*. Plant Physiol 141: 397 LP – 403

Kirchsteiger K, Pulido P, González M, Cejudo FJ (2009) NADPH thioredoxin reductase c controls the redox status of chloroplast 2-cys peroxiredoxins in *Arabidopsis thaliana*. Mol Plant 2: 298–307

König J, Baier M, Horling F, Kahmann U, Harris G, Schürmann P, Dietz K-J (2002) The plant-specific function of 2-Cys peroxiredoxin-mediated detoxification of peroxides in the redox-hierarchy of photosynthetic electron flux. Proc Natl Acad Sci 99: 5738 LP – 5743

Kono M, Terashima I (2014) Long-term and short-term responses of the photosynthetic electron transport to fluctuating light. J Photochem Photobiol B Biol 137: 89–99

Külheim C, Ågren J, Jansson S (2002) Rapid regulation of light harvesting and plant fitness in the field. Science (80-) 297: 91 LP – 93

Kumar C, Igbaria A, D’Autreaux B, Planson A-G, Junot C, Godat E, Bachhawat AK, Delaunay-Moisan A, Toledano MB (2011) Glutathione revisited: a vital function in iron metabolism and ancillary role in thiol-redox control. EMBO J 30: 2044–2056

Li X-P, Björkman O, Shih C, Grossman AR, Rosenquist M, Jansson S, Niyogi KK (2000) A pigment-binding protein essential for regulation of photosynthetic light harvesting. Nature 403: 391–395

Lima-Melo Y, Alencar VTCB, Lobo AKM, Sousa RH V, Tikkanen M, Aro E-M, Silveira JAG, Gollan PJ (2019a) Photoinhibition of photosystem i provides oxidative protection during imbalanced photosynthetic electron transport in *Arabidopsis thaliana*. Front Plant Sci 10: 916

Lima-Melo Y, Gollan PJ, Tikkanen M, Silveira JAG, Aro E-M (2019b) Consequences of photosystem-I damage and repair on photosynthesis and carbon use in *Arabidopsis thaliana*. Plant J 97: 1061–1072

Maxwell K, Johnson GN (2000) Chlorophyll fluorescence - A practical guide. J Exp Bot 51: 659–668

Mehler AH (1951) Studies on reactions of illuminated chloroplasts: I. Mechanism of the reduction of oxygen and other hill reagents. Arch Biochem Biophys 33: 65–77

Meyer AJ, Brach T, Marty L, Kreye S, Rouhier N, Jacquot J-P, Hell R (2007) Redox-sensitive GFP in Arabidopsis thaliana is a quantitative biosensor for the redox potential of the cellular glutathione redox buffer. Plant J 52: 973–986

Meyer AJ, Dick TP (2010) Fluorescent protein-based redox probes. Antioxid Redox Signal 13: 621–650

Mittler R, Vanderauwera S, Gollery M, Van Breusegem F (2004) Reactive oxygen gene network of plants. Trends Plant Sci 9: 490–498

Miyake C (2010) Alternative electron flows (water–water cycle and cyclic electron flow around psi) in photosynthesis: molecular mechanisms and physiological functions. Plant Cell Physiol 51: 1951–1963

Mizrachi A, Graff van Creveld S, Shapiro OH, Rosenwasser S, Vardi A (2019) Light-dependent single-cell heterogeneity in the chloroplast redox state regulates cell fate in a marine diatom. Elife 8: e47732

Mühlenhoff U, Molik S, Godoy JR, Uzarska MA, Richter N, Seubert A, Zhang Y, Stubbe J, Pierrel F, Herrero E, et al (2010) Cytosolic monothiol glutaredoxins function in intracellular iron sensing and trafficking via their bound iron-sulfur cluster. Cell Metab 12: 373–385

Munekage Y, Hojo M, Meurer J, Endo T, Tasaka M, Shikanai T (2002) PGR5 is involved in cyclic electron flow around photosystem I and is essential for photoprotection in Arabidopsis. Cell 110: 361–371

Nietzel T, Elsässer M, Ruberti C, Steinbeck J, Ugalde JM, Fuchs P, Wagner S, Ostermann L, Moseler A, Lemke P, et al (2019) The fluorescent protein sensor roGFP2-Orp1 monitors in vivo H_2O2_ and thiol redox integration and elucidates intracellular H2O2 dynamics during elicitor-induced oxidative burst in *Arabidopsis*. New Phytol 221: 1649–1664

Niyogi KK, Grossman AR, Björkman O (1998) Arabidopsis mutants define a central role for the xanthophyll cycle in the regulation of photosynthetic energy conversion. Plant Cell 10: 1121 LP – 1134

Ojeda V, Pérez-Ruiz JM, Cejudo FJ (2018) 2-Cys peroxiredoxins participate in the oxidation of chloroplast enzymes in the dark. Mol Plant 11: 1377–1388

Ort DR, Baker NR (2002) A photoprotective role for O_2_ as an alternative electron sink in photosynthesis. Curr Opin Plant Biol 5: 193–198

Osmond CB, Foyer CH, Bock G, Badger MR, von Caemmerer S, Ruuska S, Nakano H (2000) Electron flow to oxygen in higher plants and algae: rates and control of direct photoreduction (Mehler reaction) and rubisco oxygenase. Philos Trans R Soc London Ser B Biol Sci 355: 1433–1446

Radmer RJ, Kok B (1976) Photoreduction of O_2_ primes and replaces CO_2_ assimilation. Plant Physiol 58: 336–340

Rahantaniaina M, Tuzet A, Mhamdi A, Noctor G, Lemaire SD (2013) Missing links in understanding redox signaling via thiol / disulfide modulation□: how is glutathione oxidized in plants. 4: 1–13

Rizhsky L, Liang H, Mittler R (2003) The water-water cycle is essential for chloroplast protection in the absence of stress. J Biol Chem 278: 38921–38925

Rosenwasser S, Rot I, Meyer AJ, Feldman L, Jiang K, Friedman H (2010) A fluorometer-based method for monitoring oxidation of redox-sensitive GFP (roGFP) during development and extended dark stress. Physiol Plant 138: 493–502

Rouhier N, Couturier J, Johnson MK, Jacquot J-P (2010) Glutaredoxins: roles in iron homeostasis. Trends Biochem Sci 35: 43–52

Rouhier N, Lemaire SD, Jacquot J-P (2008) The role of glutathione in photosynthetic organisms: emerging functions for glutaredoxins and glutathionylation. Annu Rev Plant Biol 59: 143–166

Schwarzländer M, Dick TP, Meyer AJ, Morgan B (2015) Dissecting redox biology using fluorescent protein sensors. Antioxid Redox Signal 24: 680–712

Schwarzländer M, Fricker MD, Müller C, Marty L, Brach T, Novak J, Sweetlove LJ, Hell R, Meyer AJ (2008) Confocal imaging of glutathione redox potential in living plant cells. J Microsc 231: 299–316

Shikanai T (2007) Cyclic electron transport around photosystem I: genetic approaches. Annu Rev Plant Biol 58: 199–217

Shikanai T (2016) Regulatory network of proton motive force: contribution of cyclic electron transport around photosystem I. Photosynth Res 129: 253–260

Shu D-F, Wang L-Y, Duan M, Deng Y-S, Meng Q-W (2011) Antisense-mediated depletion of tomato chloroplast glutathione reductase enhances susceptibility to chilling stress. Plant Physiol Biochem 49: 1228–1237

Sonoike K (2011) Photoinhibition of photosystem I. Physiol Plant 142: 56–64

Suorsa M, Grieco M, Järvi S, Gollan PJ, Kangasjärvi S, Tikkanen M, Aro E-M (2013) PGR5 ensures photosynthetic control to safeguard photosystem I under fluctuating light conditions. Plant Signal Behav 8: e22741

Suorsa M, Jarvi S, Grieco M, Nurmi M, Pietrzykowska M, Rantala M, Kangasjarvi S, Paakkarinen V, Tikkanen M, Jansson S, et al (2012) PROTON GRADIENT REGULATION5 is essential for proper acclimation of Arabidopsis photosystem I to naturally and artificially fluctuating light conditions. Plant Cell 24: 2934–2948

Takahashi S, Badger MR (2011) Photoprotection in plants: a new light on photosystem II damage. Trends Plant Sci 16: 53–60

Takahashi S, Milward SE, Fan D-Y, Chow WS, Badger MR (2009) How does cyclic electron flow alleviate photoinhibition in *Arabidopsis*. Plant Physiol 149: 1560 LP – 1567

Terashima I, Funayama S, Sonoike K (1994) The site of photoinhibition in leaves of Cucumis sativus L. at low temperatures is photosystem I, not photosystem II. Planta 193: 300–306

Tikhonov AN (2015) Induction events and short-term regulation of electron transport in chloroplasts□: an overview. Photosynth Res 65–94

Tiwari A, Mamedov F, Grieco M, Suorsa M, Jajoo A, Styring S, Tikkanen M, Aro E-M (2016) Photodamage of iron–sulphur clusters in photosystem I induces non-photochemical energy dissipation. Nat Plants 2: 16035

Vaseghi M-J, Chibani K, Telman W, Liebthal MF, Gerken M, Schnitzer H, Mueller SM, Dietz K-J (2018) The chloroplast 2-cysteine peroxiredoxin functions as thioredoxin oxidase in redox regulation of chloroplast metabolism. Elife 7: e38194

Volpert A, Graff van Creveld S, Rosenwasser S, Vardi A (2018) Diurnal fluctuations in chloroplast GSH redox state regulate susceptibility to oxidative stress and cell fate in a bloom-forming diatom. J Phycol 54: 329–341

Waring J, Klenell M, Bechtold U, Underwood G, Baker N (2010) Waring J, Klenell M, Bechtold U, Underwood GJC, Baker NR.. Light-induced responses of oxygen photoreduction, reactive oxygen species production and scavenging in two diatom species. J Phycol 46: 1206-1217. J Phycol 46: 1206–1217

Weise SE, Weber APM, Sharkey TD (2004) Maltose is the major form of carbon exported from the chloroplast at night. Planta 218: 474–482

Wirtz W, Stitt M, Heldt HW (1982) Light activation of calvin cycle enzymes as measured in pea leaves. FEBS Lett 142: 223–226

Yamamoto H, Shikanai T (2019) PGR5-dependent cyclic electron flow protects photosystem i under fluctuating light at donor and acceptor sides. Plant Physiol 179: 588 LP – 600

Yamori W, Makino A, Shikanai T (2016) A physiological role of cyclic electron transport around photosystem I in sustaining photosynthesis under fluctuating light in rice. Sci Rep 6: 1–12

Yoshida K, Matsuoka Y, Hara S, Konno H, Hisabori T (2014) Distinct redox behaviors of chloroplast thiol enzymes and their relationships with photosynthetic electron transport in arabidopsis thaliana. Plant Cell Physiol 55: 1415–1425

Zhang Z, Jia Y, Gao H, Zhang L, Li H, Meng Q (2011) Characterization of PSI recovery after chilling-induced photoinhibition in cucumber (Cucumis sativus L.) leaves. Planta 234: 883–889

