## Supporting Appendix for "Resolving the dynamics of photosynthetically produced ROS by high-resolution monitoring of chloroplastic E_GSH_ in *Arabidopsis*"

**TABLE OF CONTENTS**

**Supp. Fig. 1:** Dose-dependent chl-roGFP2 oxidation in response to H_2_O_2._

**Supp. Fig. 2:** ETR and carbon assimilation light response curves in WT plants.

**Supp. Fig. 3:** Daily changes in ΦPSII during fluctuating light experiments in WT plants.

**Supp. Fig. 4:** Daily changes in chl-roGFP2 OxD during fluctuating light experiments in WT, *npq1* and *pgr5* plants.

**Supp. Fig. 5:** Changes in chl-roGFP2 OxD during fluctuating light cycles in WT, *npq1* and *pgr5* plants.

**Supp. Fig. 6:** Daily changes in ΦPSII during fluctuating light experiments in WT, *npq1* and *pgr5* plants.

**Supp. Fig. 7:** Daily changes in chl-roGFP2 OxD and ΦPSII during high light and recovery experiments in *gl1* plants.

**Supp. Fig. 8:** Daily changes in chl-roGFP2 OxD and ΦPSII during the third day of high-light and recovery experiments in *pgr5* plants.

**Supp. Fig. 9:** Carbon Assimilation light response curves in WT, *npq1* and *pgr5* plants.

**Supp. Fig. 10:** ETR light response curves in WT, *npq1* and *pgr5* plants.

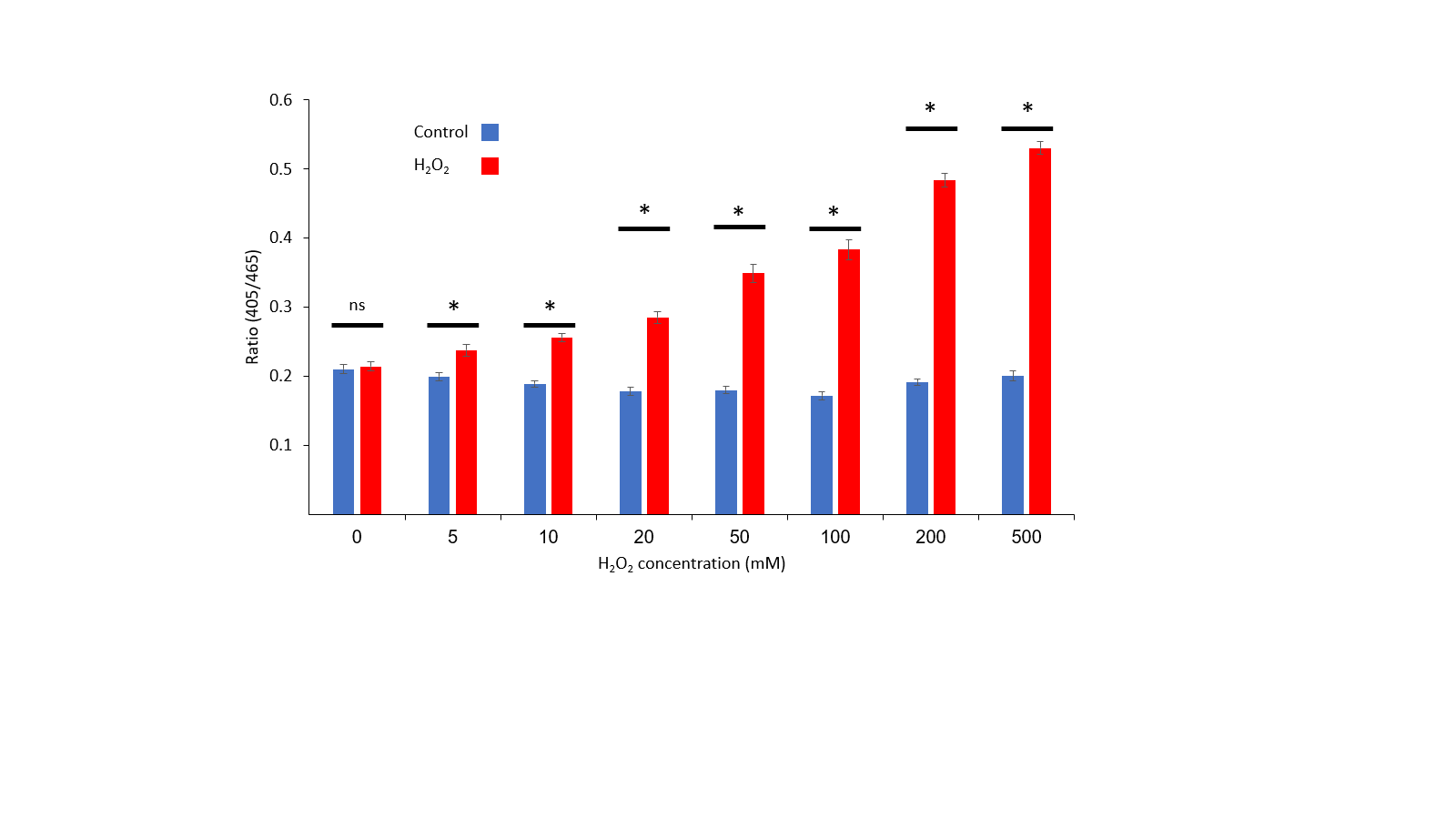

**Supp. Fig. 1. Dose-dependent chl-roGFP2 oxidation in response to H_2_O_2_.** Ratios of chl-roGFP2 emission at 510nm consequent to excitation of 405nm and 465nm (405/465) as a function of H_2_O_2_ concentration are presented. Whole-plant chl-roGFP2 fluorescence images of *Arabidopsis* plants were taken before and after treatments with increasing H_2_O_2_ concentrations (0/5/10/20/50/100/200/500mM). Ratiometric images were calculated, average ratio for each plant was extracted and values are presented as means of 24 plants ± SE. Asterisks (*) mark significant differences between untreated and H_2_O_2_ treated plants (Student t-test, P≤0.05). ns- not significant.

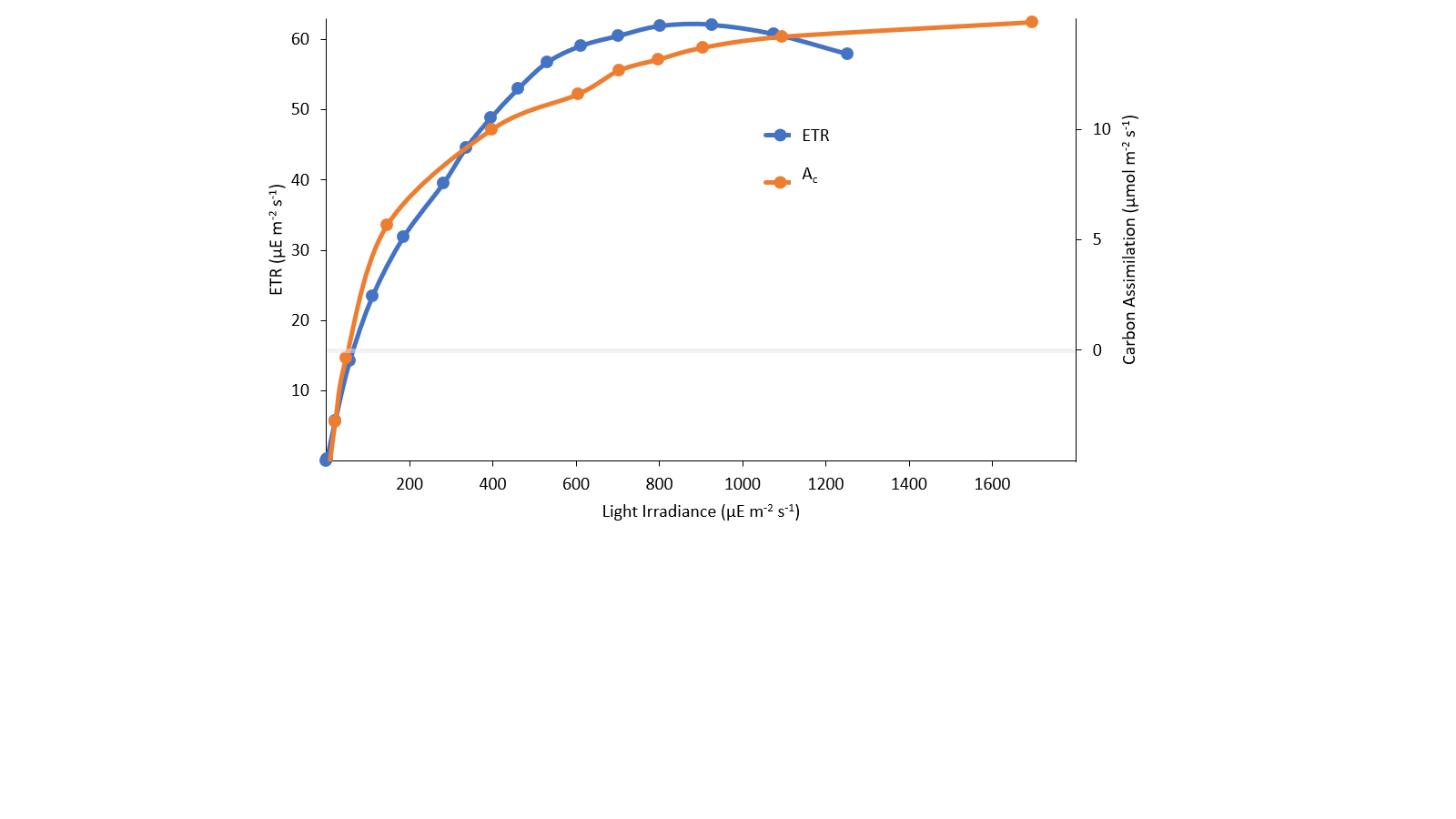

**Supp. Fig. 2. ETR and carbon assimilation light response curves in WT plants.** WT *Arabidopsis* plants were subjected to increasing light irradiances, chlorophyll fluorescence was measured and ETR was calculated (as a function of light irradiance). Values represent means of 12 plants. Carbon assimilation was calculated as a function of light irradiance using the LI-6800 system. Values were recorded from 5-6 plants grown in one pot. Total leaf area was calculated via image analysis using the Ami HT Imager and Matlab.

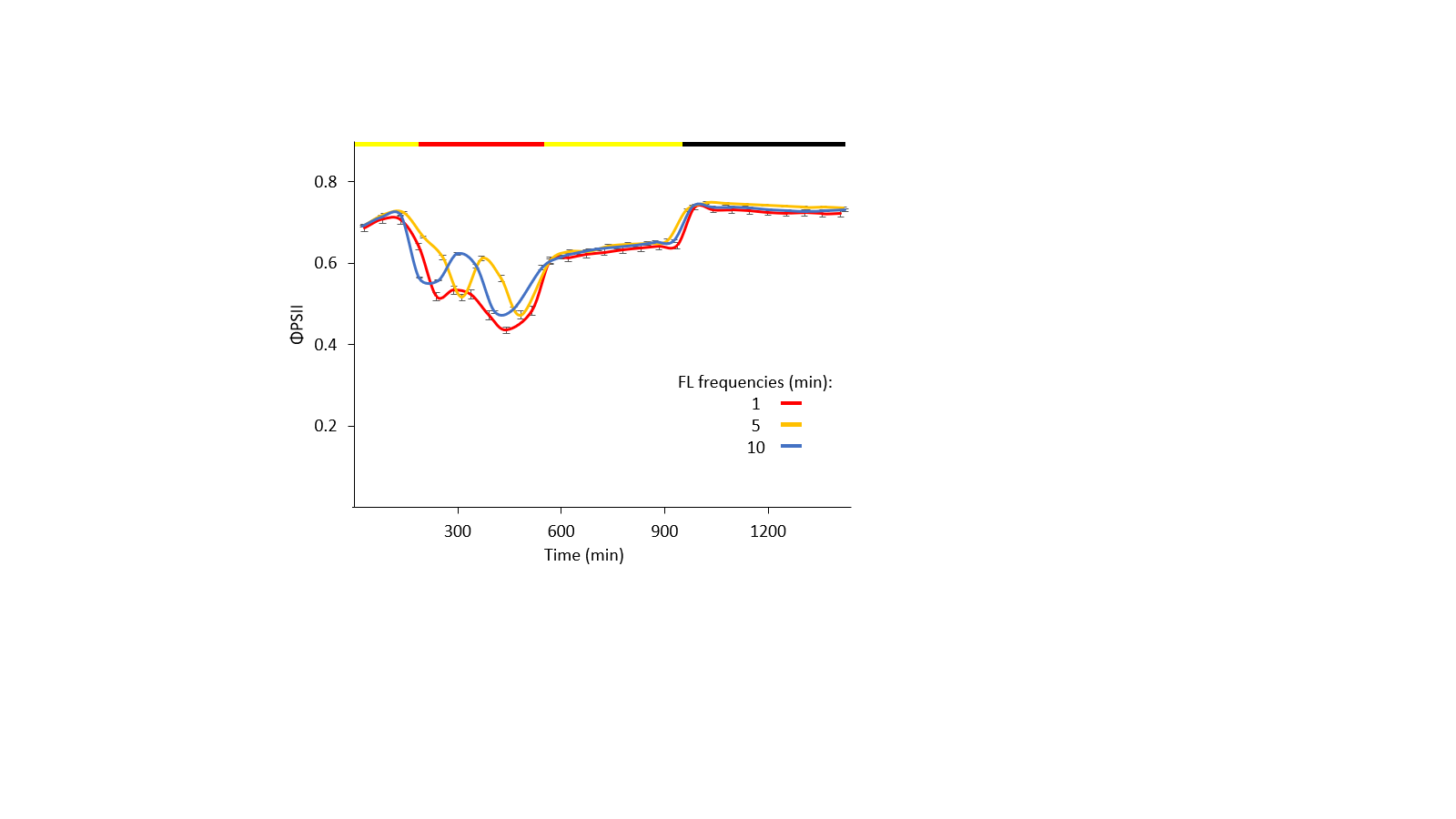
**Supp. Fig. 3.** **Daily changes in ΦPSII during fluctuating light experiments in WT plants.** WT *Arabidopsis* plants were subjected to the following light conditions during a 24-hour period: 3 hours 120µE m^-2^ s^-1^, 6 hours “fluctuating light” (between 1700µE m^-2^ s^-1^ and 120µE m^-2^ s^-1^ with a frequency of 1, 5 and 10 minutes; *i.e.* 1,5 or 10 minutes of 1700µE m^-2^ s^-1^ followed by 1, 5 or 10 minutes of 120µE m^-2^ s^-1^, respectively, and vice versa *etc*.), 7 hours 120µE m^-2^ s^-1^ and 8 hours 0µE m^-2^ s^-1^. ΦPSII was measured throughout the experiment, and values represent means (of 12 plants) ± SE as a function of the time from the experiment onset (08:30) in WT plants. The color bar denotes the light conditions: black – night, yellow – growth light, red – fluctuating light.

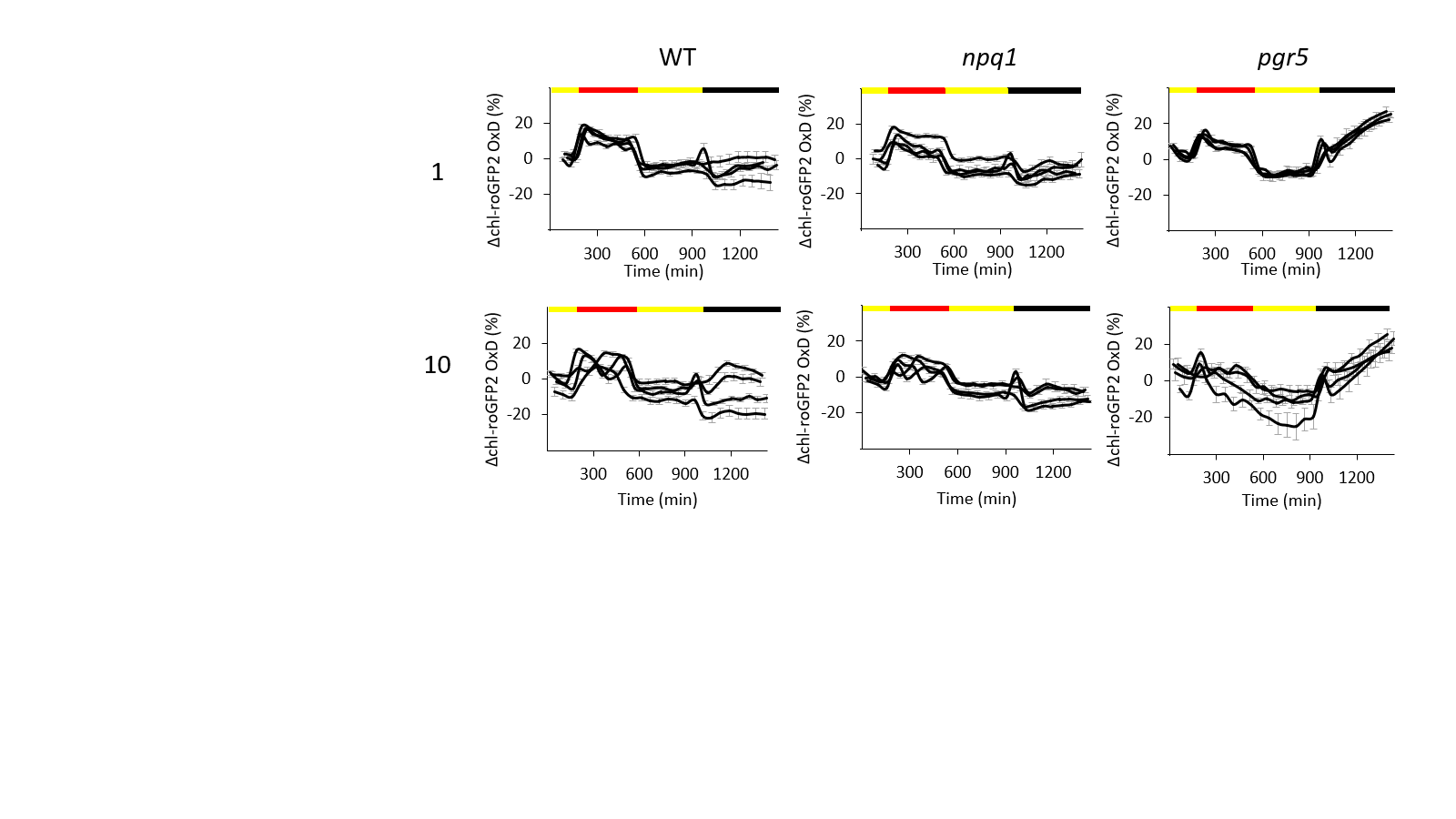

**Supp. Fig. 4. Daily changes in chl-roGFP2 OxD during fluctuating light experiments in WT, *npq1* and *pgr5* plants**. WT, *npq1* and *pgr5* *Arabidopsis* plants were subjected to the following light conditions during a 24-hour period: 3 hours 120µE m^-2^ s^-1^, 6 hours “fluctuating light” between 1700µE m^-2^ s^-1^ and 120µE m^-2^ s^-1^ with a frequency of 1 and 10 minutes (*i.e.* 1 and 10 minutes of 1700µE m^-2^ s^-1^ followed by 1 or 10 minutes of 120µE m^-2^ s^-1^, respectively, and vice versa etc.), 7 hours 120µE m^-2^ s^-1^ and 8 hours 0µE m^-2^ s^-1^. Change in chl-roGFP2 OxD relative to 'Steady state' OxD is represented as a function of the time (minutes) from the experiment onset (08:30) in WT, *npq1* and *pgr5* plants. chl-roGFP2 fluorescence was measured throughout the experiment and each line represents an independent plate consisting of 8 plants. Error bars represent standard errors. The color bar denotes the light conditions: black – night, yellow – growth light, red – fluctuating light.

**Supp. Fig. 5. Changes in chl-roGFP2 OxD during fluctuating light cycles in WT, *npq1* and *pgr5* plants.** WT, *npq1* and *pgr5* *Arabidopsis* plants were subjected to the fluctuating light experiments as described in Fig. 3. Change in chl-roGFP2 OxD relative to 'Steady state' is represented as a function of relative timing (minute) within every 1-minute frequency fluctuating light cycle (as explained in Results). The values taken from the 1700µE m^-2^ s^-1^ period within each cycle are in red box plots, while the values taken from the 120µE m^-2^ s^-1^ within each cycle are in yellow box plots, and represent means of 17-112 plants ± SE.

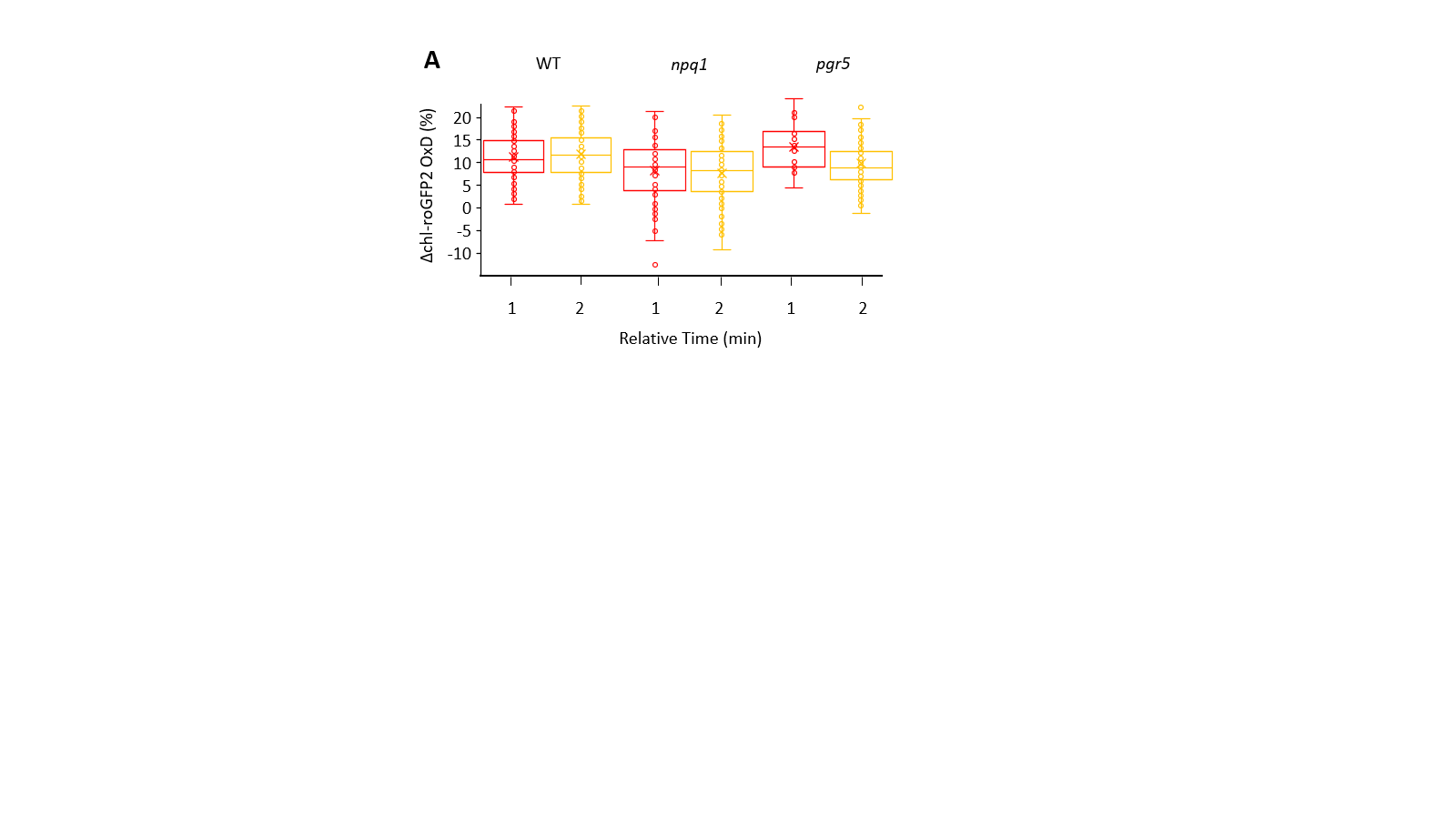

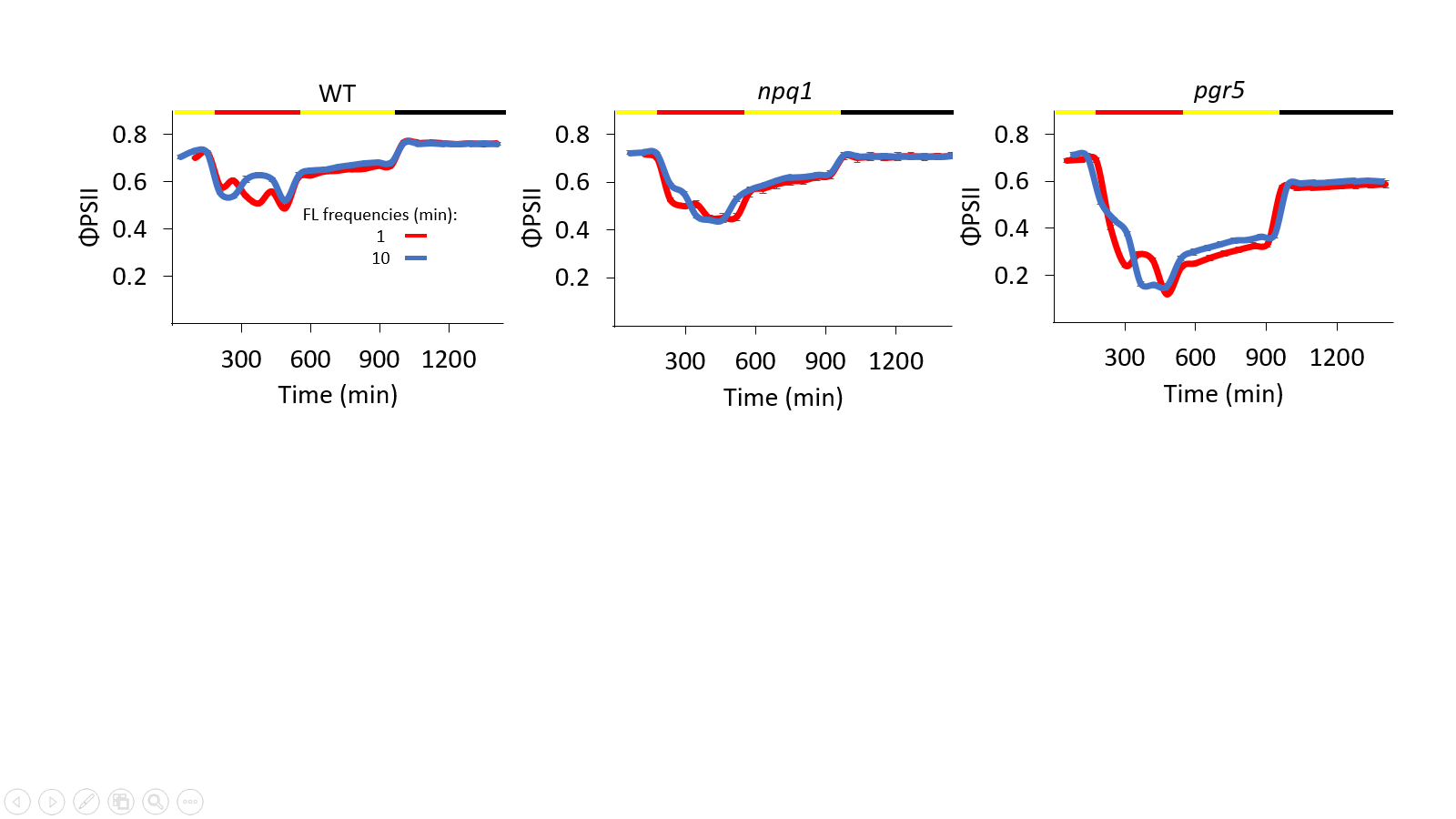

**Supp. Fig. 6. Daily changes in ΦPSII during fluctuating light experiments in WT, *npq1* and *pgr5* plants.** WT, *npq1* and *pgr5 Arabidopsis* plants were subjected to the following light conditions during a 24-hour period: 3 hours 120µE m^-2^ s^-1^, 6 hours “fluctuating light” (between 1700µE m^-2^ s^-1^ and 120µE m^-2^ s^-1^ with a frequency of 1 and 10 minutes), 7 hours 120µE m^-2^ s^-1^ and 8 hours 0µE m^-2^ s^-1^. ΦPSII as a function of the time from the experiment onset (08:30) in WT, *npq1* and *pgr5* plants. ΦPSII was measured throughout the experiment, and values represent means (of 12 plants) ± SE. The color bar denotes the light conditions: black – night, yellow – growth light, red – fluctuating light.

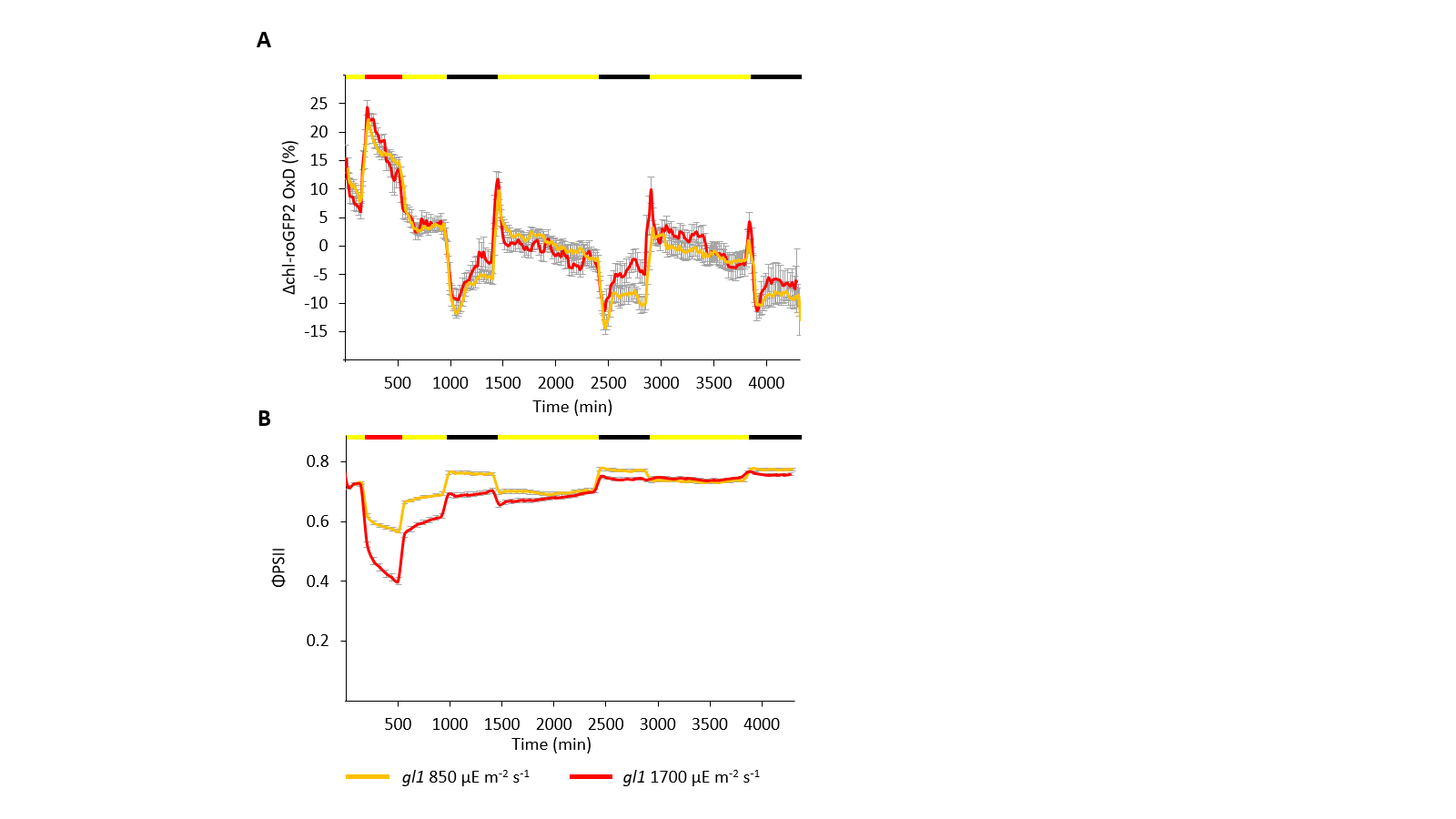

**Supp. Fig. 7.** **Daily changes in chl-roGFP2 OxD and ΦPSII during high light and recovery experiments in *gl1* plants**. *gl1 Arabidopsis* plants were subjected to the following light conditions during a 72-hour period: 3 hours 120µE m^-2^ s^-1^, 6 hours 850 or 1700µE m^-2^ s^-1^, 7 hours 120µE m^-2^ s^-1^, 8 hours 0µE m^-2^ s^-1^, followed by two 24-hour cycles of: 16 hours 120µE m^-2^ s^-1^ and 8 hours 0µE m^-2^ s^-1^. (**A**) Change in chl-roGFP2 OxD relative to steady state is represented as a function of the time (minute) from the experiment onset (08:30). For each treatment, 32 plants were used in 4 independent plates that were consolidated in a “sliding window” (n=4) display. Values represent means (of 8-32 plants) ± SE. (**B**) ΦPSII as a function of the time (minutes) from the experiment onset (08:30). ΦPSII values were derived from chlorophyll fluorescence analysis and represent means (of 12 plants) ± SE. The color bar denotes the light conditions: black – night, yellow – growth light, red – high light.

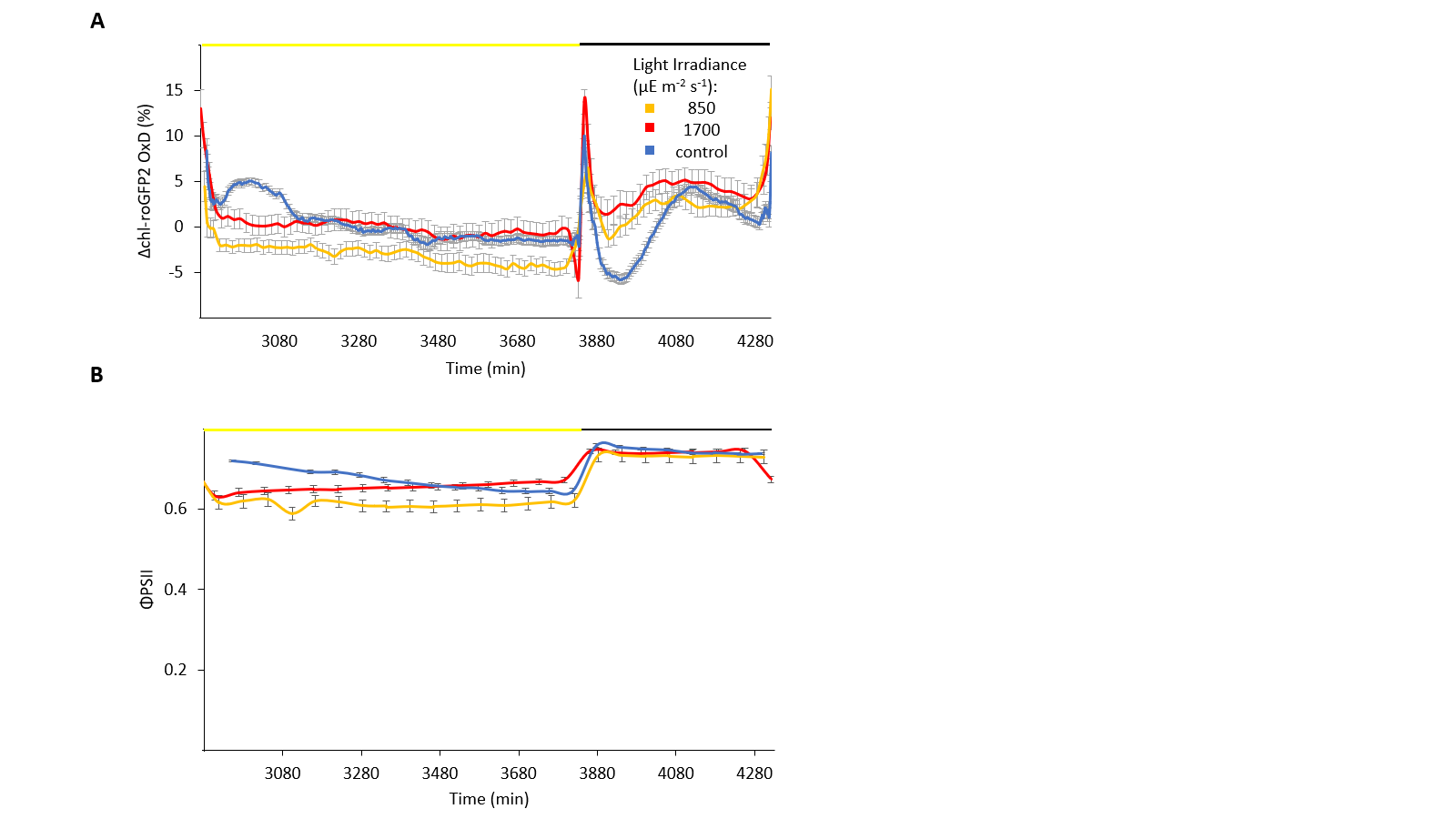

**Supp. Fig. 8. Daily changes in chl-roGFP2 OxD and ΦPSII during the third day of high-light and recovery experiments in *pgr5* plants**. *pgr5* *Arabidopsis* plants were subjected to the following light conditions during a 72-hour period: 3 hours 120µE m^-2^ s^-1^, 6 hours 850 or 1700µE m^-2^ s^-1^, 7 hours 120µE m^-2^ s^-1^, 8 hours 0µE m^-2^ s^-1^, followed by 2 24-hour cycles of: 16 hours 120µE m^-2^ s^-1^ and 8 hours 0µE m^-2^ s^-1^. (**A**)Change in chl-roGFP2 OxD relative to steady state OxD is represented during the third day of the experiment versus a “control” day as a function of the time (minutes) from the experiment onset (08:30).Values represent means (of 8 plants) ± SE and, for the control results, a "sliding window" of multiple plates (n=12) is presented as means (of 5-95 plants) ± SE. (**B**) ΦPSII during the third day of the experiment versus a “control” day as a function of the time (minutes) from the experiment onset (08:30). ΦPSII values were derived from chlorophyll fluorescence analysis and represent means (of 11 plants) ± SE. The color bar denotes the light conditions: black – night, yellow – growth light, red – high light.

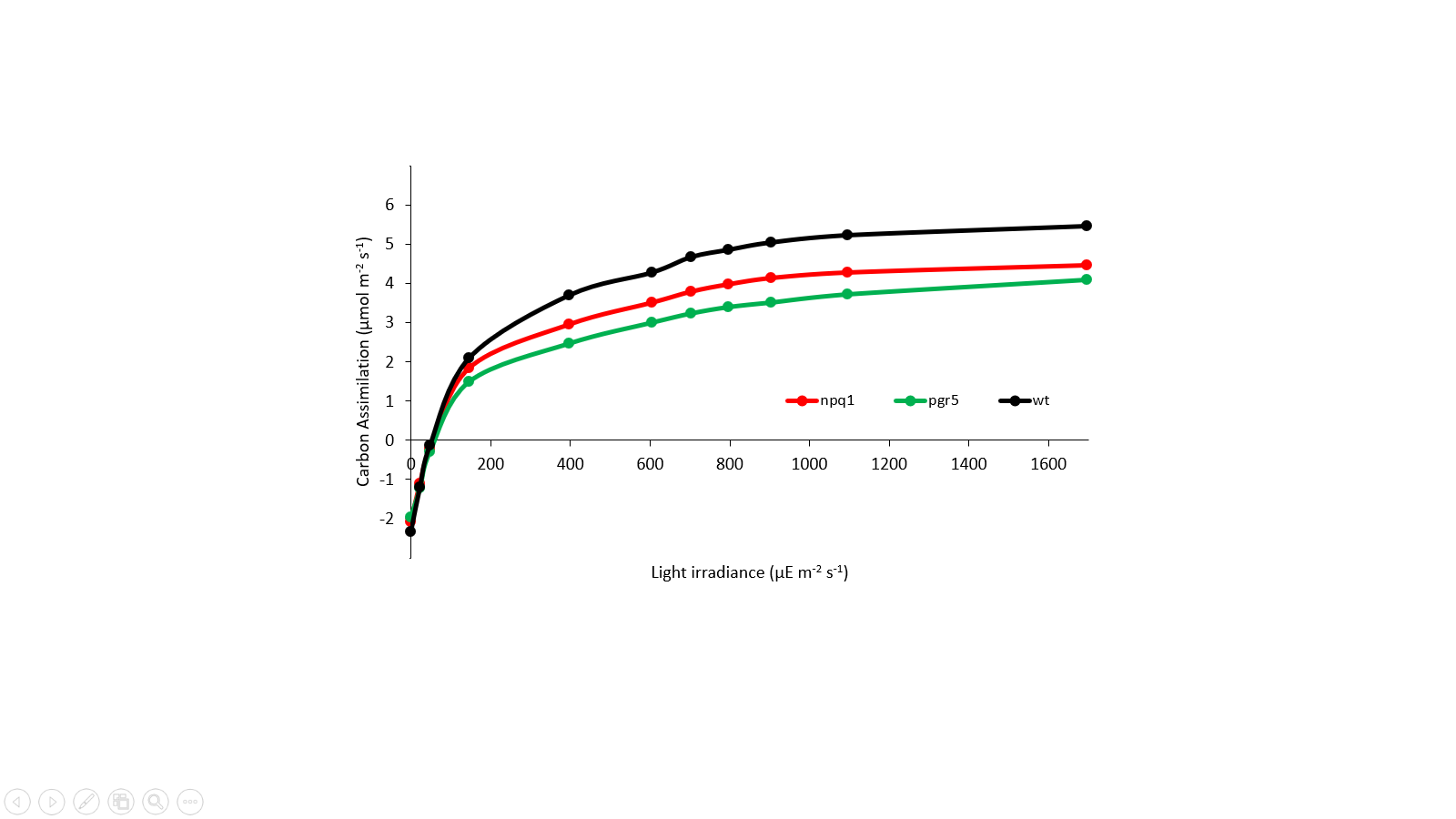

**Supp. Fig. 9. Carbon Assimilation light response curves in WT, *npq1* and *pgr5* plants.** WT, *npq1* and *pgr5* *Arabidopsis* plants were subjected to increasing light irradiances. Carbon Assimilation was calculated as a function of light irradiance using the LI-6800 system (Values represent means of 5-6 plants).

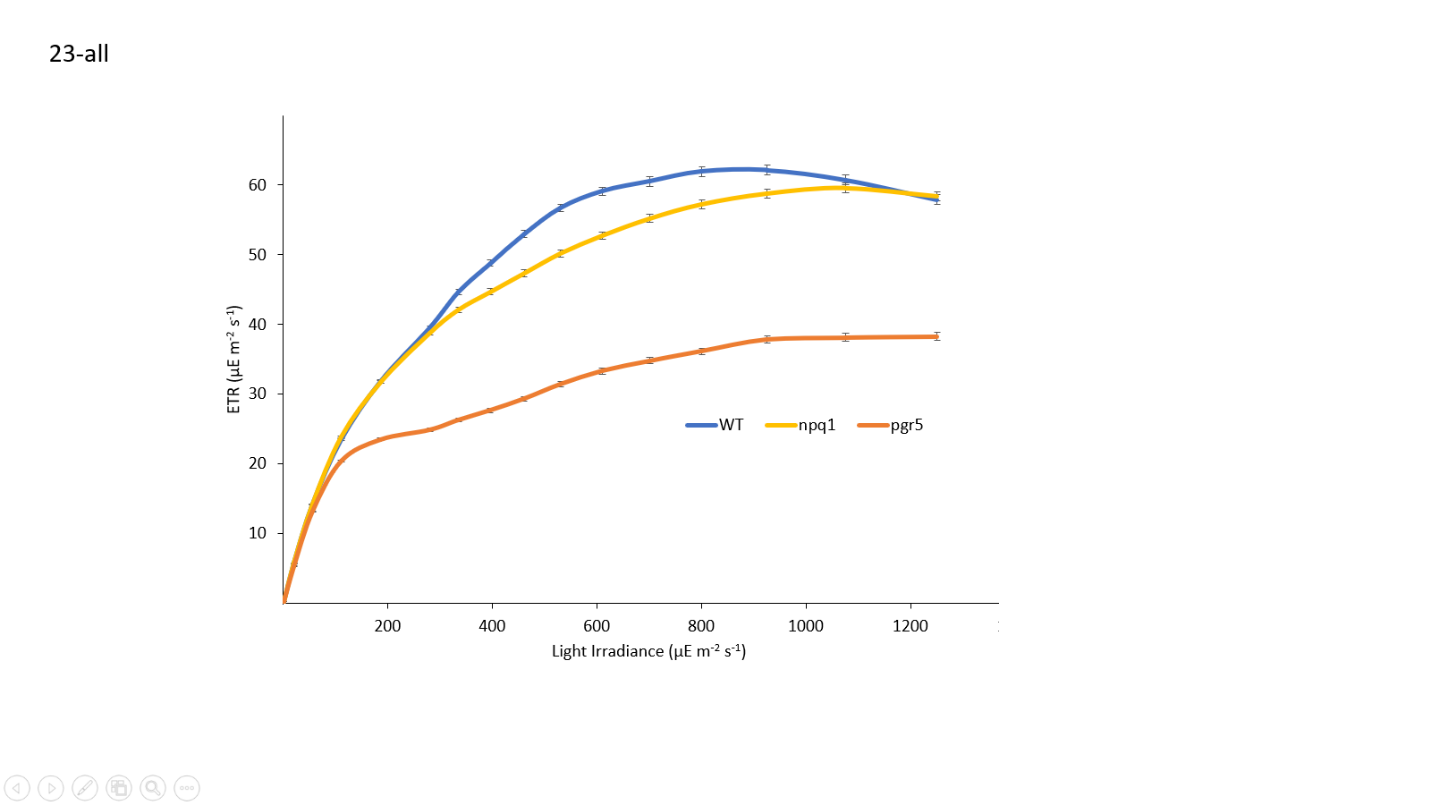

**Supp. Fig. 10. ETR light response curves in WT, *npq1* and *pgr5* plants.** WT, *npq1* and *pgr5* *Arabidopsis* plants were subjected to increasing light irradiances. Chlorophyll Fluorescence was measured and ETR was calculated as a function of light irradiance (See Methods, Values represent means of 12 plants ± SE).
